# Opaque Ontology: Neuroimaging Classification of ICD-10 Diagnostic Groups in the UK Biobank

**DOI:** 10.1101/2024.04.15.589555

**Authors:** Ty Easley, Xiaoke Luo, Kayla Hannon, Petra Lenzini, Janine Bijsterbosch

**Affiliations:** Department of Radiology, Washington University School of Medicine, Saint Louis, Missouri 63110, USA

**Keywords:** UK Biobank, machine learning, neuroimaging, mental health disorders, nervous system diseases

## Abstract

**Background:** The use of machine learning to classify diagnostic cases versus controls defined based on diagnostic ontologies such as the ICD-10 from neuroimaging features is now commonplace across a wide range of diagnostic fields. However, transdiagnostic comparisons of such classifications are lacking. Such transdiagnostic comparisons are important to establish the specificity of classification models, set benchmarks, and assess the value of diagnostic ontologies.

**Results:** We investigated case-control classification accuracy in 17 different ICD-10 diagnostic groups from Chapter V (mental and behavioral disorders) and Chapter VI (diseases of the nervous system) using data from the UK Biobank. Classification models were trained using either neuroimaging (structural or functional brain MRI feature sets) or socio-demographic features. Random forest classification models were adopted using rigorous shuffle splits to estimate stability as well as accuracy of case-control classifications. Diagnostic classification accuracies were benchmarked against age classification (oldest versus youngest) from the same feature sets and against additional classifier types (K-nearest neighbors and linear support vector machine). In contrast to age classification accuracy, which was high for all feature sets, few ICD-10 diagnostic groups were classified significantly above chance (namely, demyelinating diseases based on structural neuroimaging features, and depression based on socio-demographic and functional neuroimaging features).

**Conclusion:** These findings highlight challenges with the current disease classification system, leading us to recommend caution with the use of ICD-10 diagnostic groups as target labels in brain-based disease prediction studies.

## Background

Many studies have trained machine learning classifiers on features derived from non-invasive structural and/or functional neuroimaging data to differentiate between cases and healthy controls in a range of diseases (for a review see [1]). In such disease classification studies, the definition of cases and controls is commonly based on standardized disease ontologies such as the ICD-10 or the DSM-V and assessed via structured clinical interviews and/or health records. However, the vast majority of published diagnostic classification efforts are single-disease studies performed in disease-specific cohorts. As such, a comprehensive analysis across diseases in population data is lacking. The goal of this study is to leverage epidemiological data to comprehensively assess the ability to accurately classify 17 ICD-10 diagnostic groups from chapters V (mental and behavioral disorders) and VI (diseases of the nervous system) based on neuroimaging features.

The UK Biobank (UKB) offers the first available neuroimaging dataset that adopts an epidemiological approach in terms of its prospective recruitment strategy and large sample size [2]. At the time of writing, neuroimaging data for tens of thousands of participants recruited from the National Health Service (NHS) database had been acquired and released, with data acquisition still ongoing towards the goal of N=100,000 [3]. Although the UKB has some healthy volunteer selection bias resulting from the opt-in choice of participation [4], there are no explicit health-based inclusion/exclusion criteria apart from standard MRI contraindications for neuroimaging. As such, participants with a wide range of ICD-10 diagnoses (derived from clinical records) are included in the UKB cohort [5]. This work uses the UKB dataset to systematically compare brain-based classification models across 17 different ICD-10 diagnostic groups.

This study provides a unique lens on brain-based classifications across a large set of mental and neurological ICD-10 diagnostic groups. There are a number of reasons why such transdiagnostic comparisons of diagnostic classification models are important. First, such comparisons are needed to assess the specificity of classification models (in addition to their accuracy/sensitivity). Machine learning tools are increasingly adopted in clinical care settings, yet their disease-specificity cannot be assessed in single-disease studies. Although some transdiagnostic comparisons have been performed across related disorders (e.g., bipolar-schizophrenia [6]; mild cognitive impairment-Alzheimer’s disease [7]), broader comparisons are lacking. Second, our findings provide a comprehensive benchmark for future research on diagnostic classification models in the UKB and beyond. The UKB cohort will become an increasingly valuable resource to train disease classification models as it follows participants longitudinally, capturing hospital and death records. As such, our findings provide an important benchmark for future efforts. Third, our findings shed light on the limited validity of the ICD-10 diagnostic ontology, consistent with other research pointing to the limited reliability of diagnostic coding systems including the ICD-10 [8] and DSM-V [9, 10]. Importantly, the use of suboptimal clinical labels as targets for machine learning models impedes meaningful biomarker discovery [11, 12].

In total, this work included N=5,861 unique cases and the same number of carefully matched healthy controls. We trained and tested over 400 diagnostic classification models to gather a comprehensive overview of results. Despite well-matched samples of moderate to large sample sizes (N range: 250 - 2,658), mean classification accuracies ranged from chance (0.5) up to 0.69, and many diagnostic groups could not be classified significantly beyond chance. As such, our comprehensive results revealed limits of the ICD-10 ontology (as available in the UKB) and provide an important benchmark for future work in the UKB and beyond.

## Data Description

### 1. Case selection

We used UKB variable ID 41270 to collect ICD-10 information for all participants. We focused on ICD-10 diagnostic groups in Chapter V (Mental and behavioral disorders) and Chapter VI (Diseases of the nervous system) because these diagnostic groups are most relevant for brain-based classification. The number of UKB participants with complete neuroimaging data at each of the first levels of the ICD-10 hierarchy (e.g., F00-F09; which we term broad diagnostic groups) and at each of the second levels of the ICD-10 hierarchy (e.g., F00 separately; which we term narrow diagnostic groups) were determined. ICD-10 groups with N≥125 cases at the narrow diagnostic category level were retained. If no individual narrow diagnostic category contained N≥125, but the combined broad diagnostic category did contain N≥125, then the broad diagnostic category was retained. As a result, we retained 17 ICD-10 diagnostic groups (see Table S1). We also selected a unique case list by removing all subjects who were in more than 1 of the 17 diagnostic groups (Table S2). This unique case list was used for multiclass classification.

### 2. Matched healthy controls

For each of the 17 diagnostic groups, we matched cases to controls to achieve 17 fully balanced case-control groups for classification. The total number of UKB participants with complete neuroimaging data but with no ICD-10 labels in either Chapter V (Mental and behavioral disorders; all F classes) or Chapter VI (Diseases of the nervous system; all G classes) was N=31,225, which composed our pool of healthy controls. Out of this pool, controls were selected for each case to match sex, age, and resting state head motion as closely as possible (in this order of priority). Each case (combined N=5861) was matched to a unique control participant across all combined diagnostic groups. The matching procedure resulted in perfectly matched groups for sex (*χ*^2^ *p*=1 for all diagnostic groups) and no significant group differences for age (p>0.3) nor for head motion (p>0.7).

### 3. Matched sample size subgroups

The resulting ICD-10 diagnostic groups varied substantially in sample size (ranging from 125 - 1,329 cases). To test whether classification performance was impacted by sample size, we repeated analyses after subsampling each ICD-10 diagnostic group to match the minimum sample size (125 cases + 125 controls). Subsampling was performed by matching the cases from each ICD-10 diagnostic group to the ICD-10 diagnostic group with the minimum sample size (G35-G37; Demyelinating diseases of the central nervous systems) for sex, age, and resting state head motion using the same procedure described in section 2 above. This subsampling procedure therefore additionally removed potential differences in confounding variables between ICD-10 diagnostic groups.

## Analyses

Random forest classification models were trained separately for each diagnostic category across 100 shuffle split repeats with 80% training data and 20% validation data (see Table S3). The primary neuroimaging features used to drive the classification algorithms included two sets of structural measures (285 surface-based measures, or 153 volumetric measures; Table S4). Significance testing was performed by comparing the resulting 100 classification accuracies for each shuffle split against chance level (0.5) as described in Methods section 6. Follow-up analyses were performed to test whether the primary classification results could be improved upon by performing: i) classifications using matched sample sizes, ii) classifications using a multiclass algorithm, iii) alternative classification models (support vector and k-nearest neighbors classifiers), and iv) alternative features sets (from functional neuroimaging or sociodemographic information; see Table S5). Furthermore, diagnostic classification accuracies were benchmarked based on age classification by differentiating between extreme ends of the age distribution (youngest versus oldest). Two age classification groups were generated that matched the sample sizes of the largest and smallest ICD-10 diagnostic groups. A more detailed description of the analytical approaches can be found in the Methods section.

### 1. Diagnostic classification using structural neuroimaging features

Only the “Demyelinating diseases” ICD-10 diagnostic group (G35-37) was classified significantly above chance by both surface-based structural neuroimaging features (*μ_acc_* = 0.63, *p* = 0.013, n=248; Fig. 1 & Table S6) and volumetric features (*μ_acc_* = 0.68, *p* = 0.0094, n=248; Fig. S1 & Table S6) after false discovery rate correction over the two structural feature sets (volume and surface). Structural neuroimaging data features failed to classify the remaining 16 diagnostic groups significantly more accurately than chance (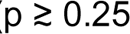; Fig. 1).

**Figure 1.**
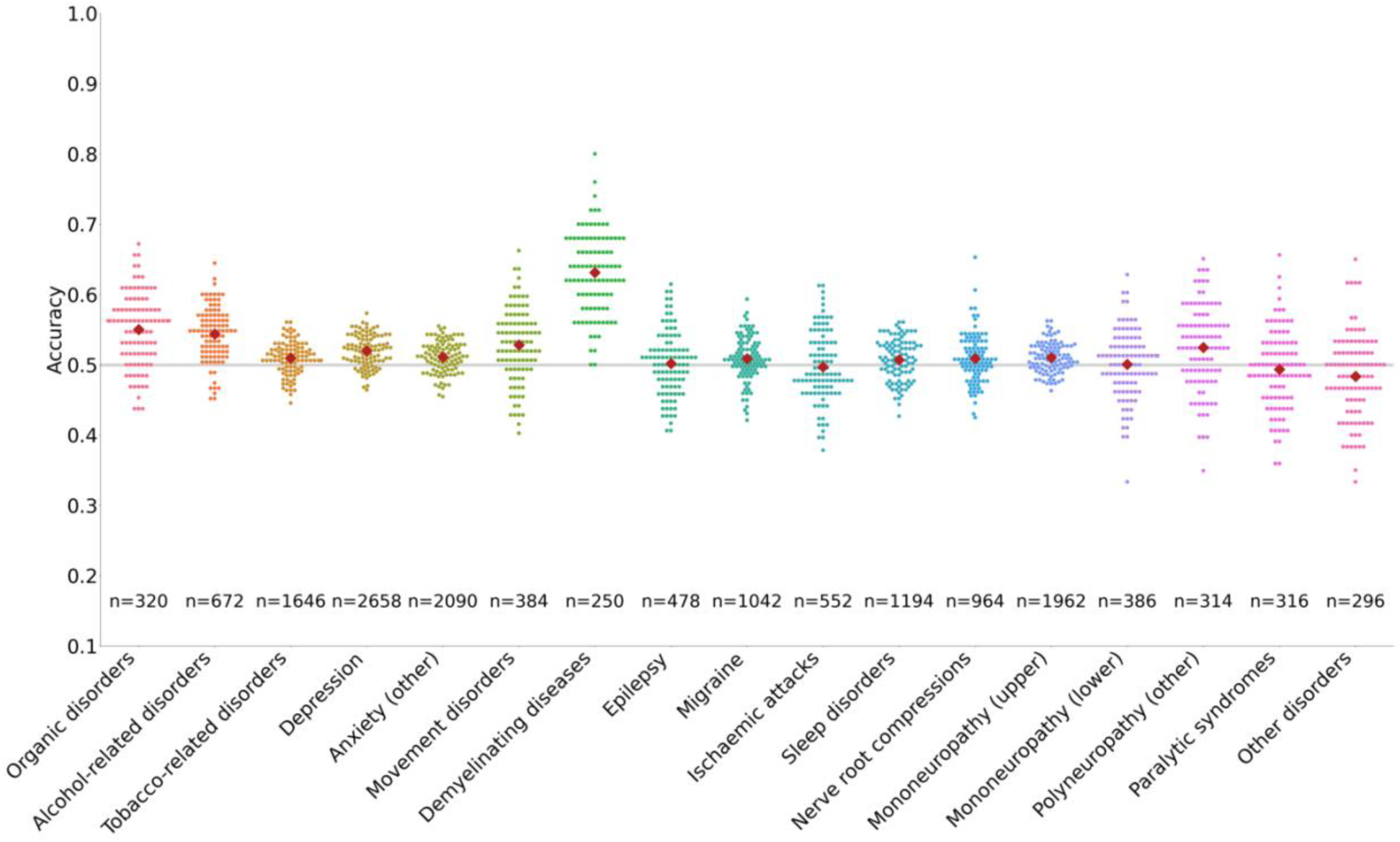
Diagnostic classification based on surface-based structural neuroimaging features. Classification accuracy distributions across ICD-10 diagnostic groups for cortical surface features derived from T1-weighted structural MRI data. Only demyelinating diseases were classified significantly above chance. Mean classification accuracy across splits is shown as a single red dot within each distribution. Matching results using the volumetric feature sets are available in Fig. S1 and all numeric results are available in Table S6.

### 2. Diagnostic classification corrected for sample size

To mitigate the impact of varying sample sizes, we repeated the diagnostic classification after subsampling each ICD-10 diagnostic group to match the minimum sample size of 125 cases and 125 matched controls. With a uniform size of 125 for diagnostic classification, the main results shown in Fig. 1 were replicated. Namely, only the “Demyelinating diseases” ICD-10 diagnostic group (G35-37) was classified significantly above chance by surface-based structural neuroimaging features(μ_*acc*_ =0.64,*p*=0.01, n=125; Fig 2a). Full numeric results for surface-based and volumetric features sets are available in Table S7.

**Figure 2.**
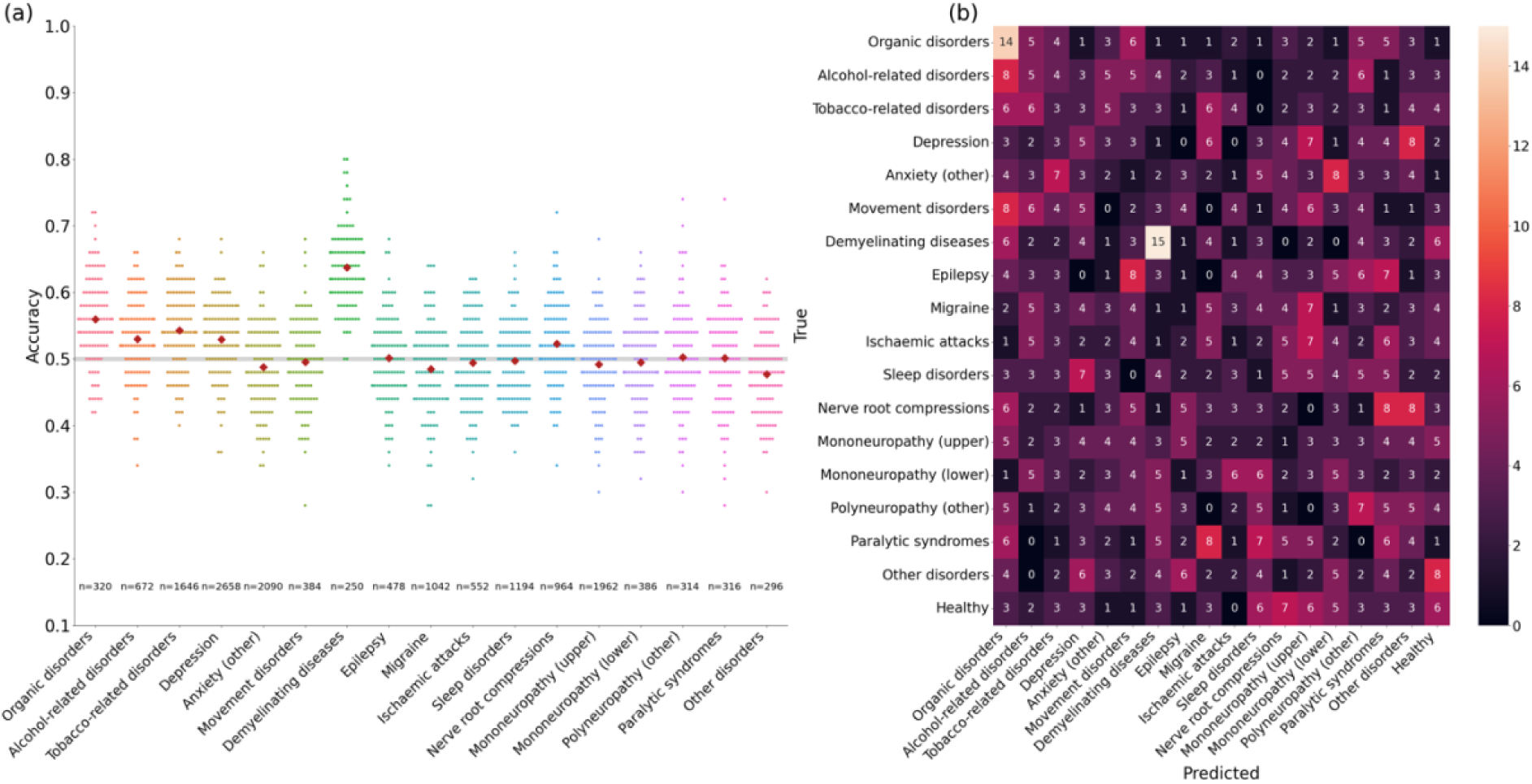
(a) Size-corrected diagnostic classification based on surface-based structural neuroimaging features. Classification accuracy distributions across ICD-10 diagnostic groups matched in sample size. All classification results matched for sample size are reported in Table S7. (b) Size-corrected multiclass diagnostic classification. Confusion matrix of the multiclass classification for all 17 ICD-10 diagnostic groups along with healthy controls.

### 3. Multiclass classification of ICD-10 diagnostic groups

To test whether our diagnostic classifications may improve by combining all diagnoses into one model, we trained a multiclass classifier. Participants with multiple diagnostic labels were removed to ensure disjoint groups and sample sizes were matched across ICD-10 diagnostic groups to avoid the impact of unbalanced groups (see Methods for more details; Table S2). Multiclass classification results are summarized in the confusion matrix in Fig. 2B. With a uniform sample size of N=59 per ICD-10 diagnostic group for multiclass classification, 92% (977 off diagonal individuals out of N=1,062) of the subjects were misclassified across all the diagnostic groups. Aligning with the results from diagnostic classification, the ICD-10 diagnostic group with the highest number of correct classifications was “Demyelinating diseases”. Notably, although the individual group sizes are small due to the removal of participants with multiple diagnostic labels, the total sample size for the multiclass classification algorithm remained relatively large (N=1,062). Comparable results using the volumetric feature set can be found in Fig. S2.

### 4. Comparison against alternative classification models

We also classified ICD-10 diagnostic groups from structural features using support vector classification and *k*-nearest neighbors classification to check the robustness of our findings in other classification paradigms. The results were similar across classification models and the two additional models did not significantly predict any additional diagnostic groups beyond the random forest classification results (Fig. 3; Table S8).

**Figure 3.**
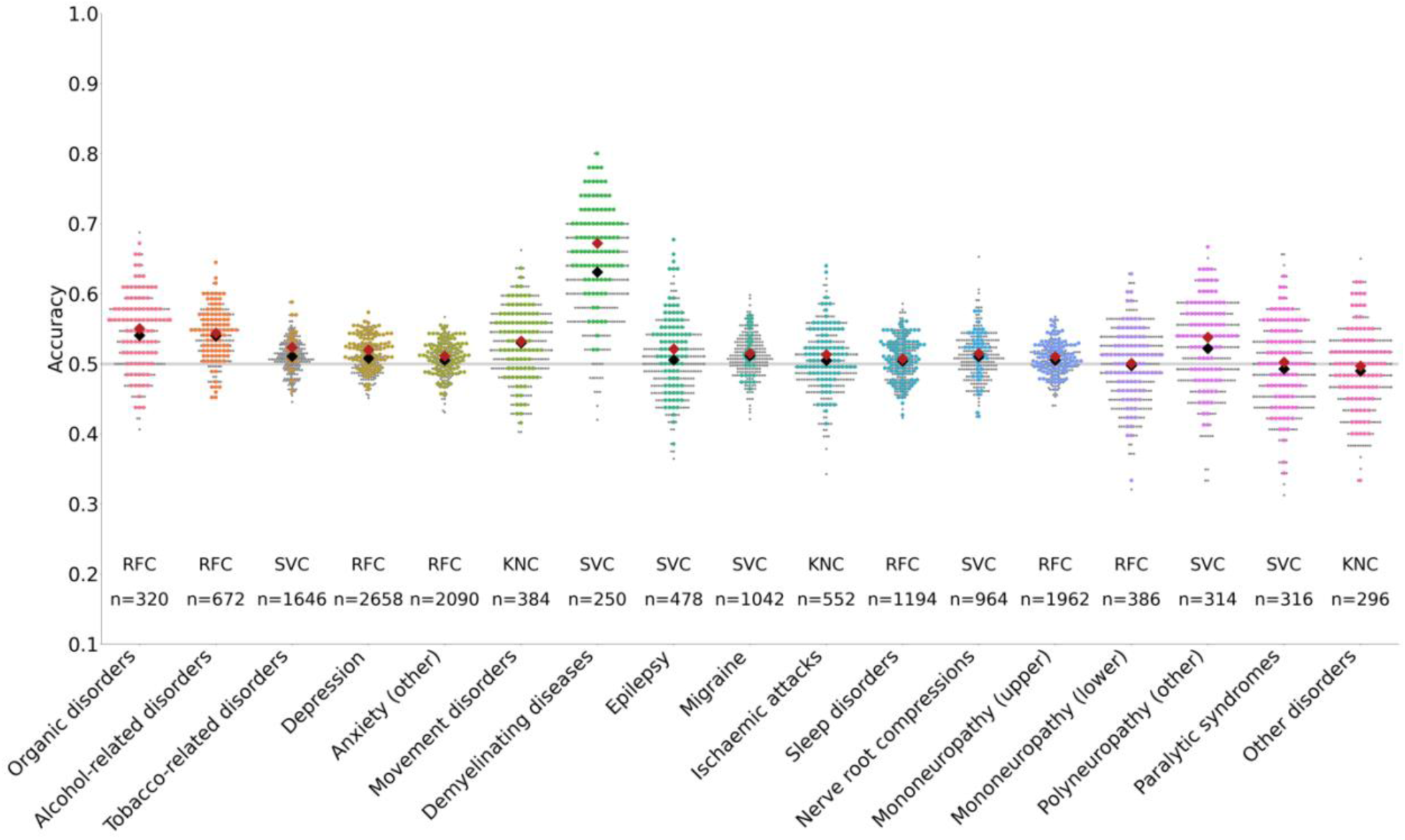
Effect of classification models. Classifier performance for all ICD-10 diagnostic groups on T1-weighted structural MRI surface data. Results for all classification models are shown in gray dots and results for the best performing classification model are shown in colored dots. Because classification accuracy distributions for different classifiers heavily overlap, few gray dots are visible. Large red diamonds show mean classification accuracy of the best-performing classifier, while large black diamonds show mean classification accuracy across all classifiers. All classification results across different models are reported in Table S8.

### 5. Comparison against alternative feature sets

We furthermore tested whether adopting alternative feature sets may improve the classification accuracy of ICD-10 diagnostic groups. Specifically, we tested 17 different feature sets derived from resting state functional MRI (see Supplementary Materials section 3) and one sociodemographic features set (based on [13], see Supplementary Materials section 4). Within each diagnostic group, multiple comparisons correction using false discovery rate control was performed across the total number of feature sets tested (i.e., across 20 feature sets, including the 2 primary structural MRI feature sets, 17 functional MRI feature sets, and 1 sociodemographic feature set). Because the multiple comparisons correction was more stringent than when using only structural features, demyelinating diseases (G35-37) were not significantly classified after correcting across all feature types. Instead, we found that the depression group (F32) was classified significantly above chance after correction across multiple feature groups (Fig. 4). Depression classification accuracy was highest when using the sociodemographic feature set (*μ_acc_*=0.58, *p*=3.5e-3, n=2692), but was also classified significantly above chance by several resting state feature sets after multiple comparisons correction (see Table S9). Within the 16 diagnostic groups that were not classified significantly above chance, we found little variation in classification accuracy across feature sets. This suggests that classification accuracy was generally low and largely insensitive to feature choice, given the statistical power available in this study.

**Figure 4.**
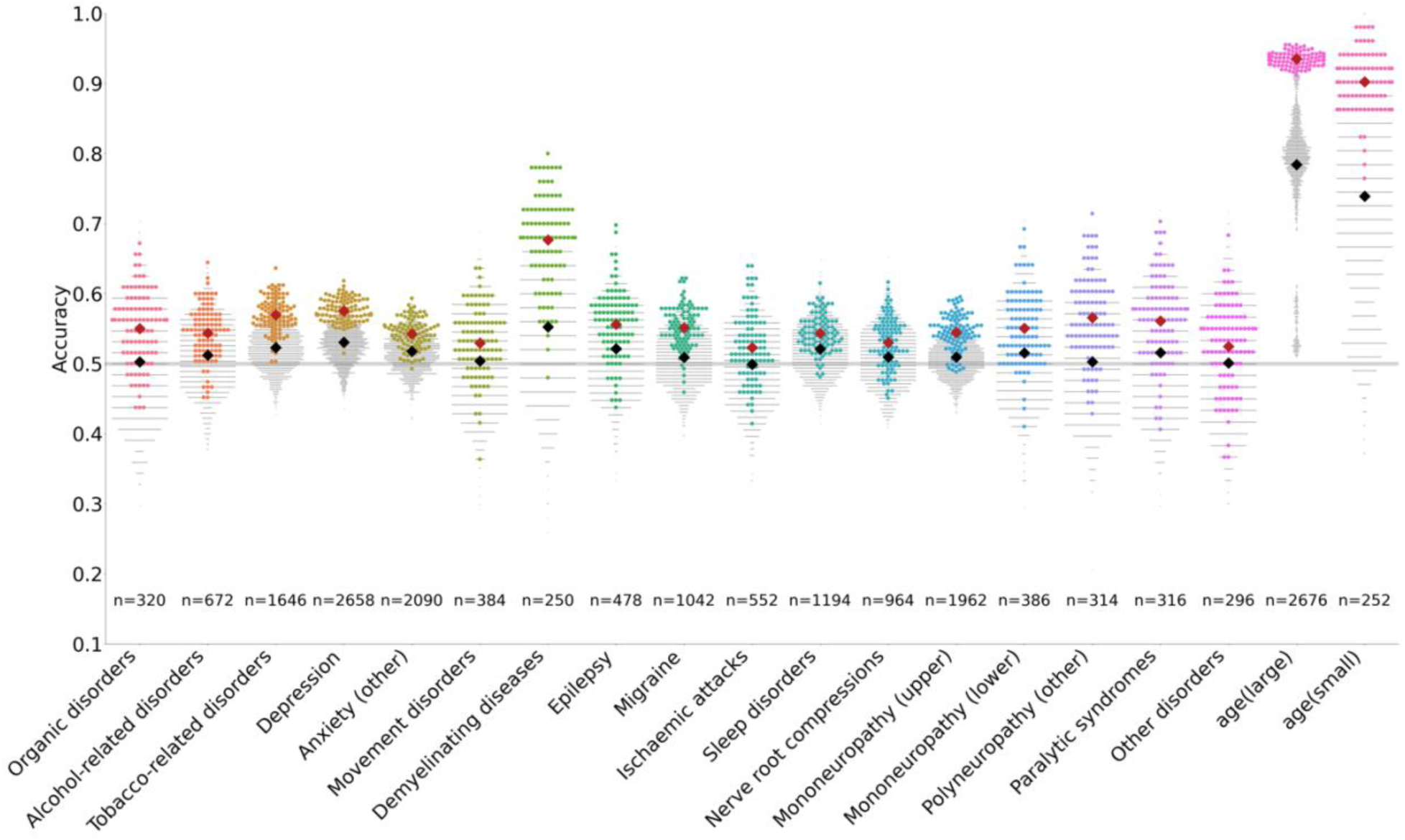
Diagnostic and age classification across feature sets. Summary of classification accuracy distributions across all 17 diagnostic groups for structural, functional, and sociodemographic classification features. In each classification group’s swarm plot, the accuracy distribution of the feature set with the highest mean accuracy is shown in color; the rest are shown in gray. Of the diagnostic groups, only the depression group (N=2,692) was classified significantly above chance after multiple comparison corrections. Depression was significantly classified by sociodemographic features and three functional MRI features (full network connectivity matrices for the Schaefer parcellation, full network connectivity matrices for ICA parcellation at rank 150, and at rank and 300). Benchmark results for age (youngest versus oldest) classifications in both a large and small group (size-matched, respectively, to the largest and smallest diagnostic groups) are shown on the right, and approximate an upper bound on diagnostic classification accuracy in terms of effect size vs. sample size. All classification results are reported in Table S9.

### 6. Comparison against age classification

To establish a plausible upper bound on classification accuracy of our random forest classification model and neuroimaging feature sets, we classified age (oldest vs. youngest; see Methods section 10). Specifically, we classified age (oldest vs. youngest) in both a large group, size - matched to the depression group (N=2,676; our largest ICD-10 diagnostic group; F32) and a small group, size-matched to the demyelinating diseases group (N=250; our smallest ICD-10 diagnostic group; G35-37). Structural neuroimaging features (derived from surface data) best classified age in both the small (*μ_acc_*=0.90, *p*=1.3e-15, n=246) and large (*μ_acc_*=0.94, *p*=1.4e-66, n=2676) groups. All considered features classified age significantly above chance in the large age group (*p* < 1.6e-3; Fig. 4). Nearly all features were able to significantly classify age above chance in the small group as well (*p* < 1.8e-3), with the exception of partial connectivity matrices derived from the Schaefer parcellation (*p*=0.35) and independent component analyses of ranks 150 (*p*=0.085) and 300 (*p*=0.34). See Tables S9 for a detailed summary of results. These results confirm that the classification model and neuroimaging feature sets adopted in this work can achieve higher classification accuracies than those observed across all ICD-10 diagnostic groups.

## Discussion

The goal of this study was to systematically compare brain-based classification models across 17 different ICD-10 diagnostic groups. Our findings revealed that most ICD-10 diagnostic groups were not classified significantly above chance from neuroimaging or sociodemographic data in the UK Biobank. We found that only a single diagnostic group (“Demyelinating diseases”, G35-37) could be accurately classified from structural neuroimaging features alone. After adding additional feature sets, the largest group (“Depression”, F32) was classified significantly above chance by both sociodemographic features and several functional neuroimaging features, but not by structural neuroimaging features. Size-matched classification groups and a multiclass formulation of the problem both yielded similar insights. Classifications with random forest, support vector, and *k*-nearest neighbor classifiers all gave comparable results. Ultimately, no set of either neuroimaging or sociodemographic features was able to classify more than one ICD-10 diagnostic group significantly above chance. By contrast, nearly all neuroimaging features were able to classify age with high accuracy in samples that were size-matched to both our smallest and largest diagnostic groups.

Paradoxically, the two diagnostic groups that were significantly classified by one or more feature sets represented the largest group (depression; N=2,658) and the smallest group (demyelinating diseases, N=250) included in this work. These results are consistent with the notion that the statistical power to detect an effect depends on three factors: i) the magnitude of the effect of interest in the population, ii) the sample size used to detect the effect, and iii) the statistical significance criterion used in the test. The demyelinating diseases diagnostic group revealed outsized sensitivity, suggesting a relatively larger magnitude of effect of interest in this diagnostic population. The impact of the statistical significance criterion, and particularly the multiple comparisons burden, can be observed in the finding that the demyelinating disease group was significantly classified from our primary structural neuroimaging feature sets (controlling across 2 results), but this result no longer reached significance after correction across the expanded number of feature sets (controlling across 20 results; Table S9). These results highlight the importance of balancing hypothesis-driven choices with data-driven exploration. Notably, our statistical significance criterion was relatively strict by using the standard deviation - rather than standard error of the mean - as the variance criterion for division. Although a larger number of results would reach significance under a more lenient significance criterion, this does not take away from the central interpretation that classification accuracy was close to chance for the majority of ICD-10 diagnostic groups.

For the depression group, our findings revealed that classification accuracy based on functional neuroimaging features sets outperformed classification accuracy based on structural neuroimaging feature sets (Table S9). These findings are consistent with the conceptualization of depression as driven by circuit dysconnectivity [15, 16]. Although structural brain correlates of depression have also routinely been reported in the literature [17, 18], our findings are partially consistent with a recent meta analysis that did not find significant convergence of structural or functional brain correlates of depression [19]. Importantly, the individual studies included in this meta analysis were relatively underpowered (N<100), further emphasizing the need for more well-powered research [14]. It is possible that our findings may be linked to the larger feature space for functional features (>10,000 features compared to <300 structural features), although our rigorous shuffle split validation framework protects against overfitting. Overall, our findings may suggest that functional neuroimaging features are more sensitive to depression than structural neuroimaging features, although future research is needed to confirm this finding.

Our classification results showed little variation in performance over different classifiers, classification features, and target diagnostic groups. Sociodemographic features were also unable to accurately classify ICD-10 diagnostic groups, despite previous work showing higher classification accuracy of sociodemographic features for other phenotypes [13]. In contrast to the high accuracy of age classification across all feature types and observed sample sizes, we found ICD-10 diagnostic groups to have nearly uniformly low classification accuracy. Our investigation across feature types, classification algorithms, and classification targets therefore suggests that ICD-10 diagnostic groups may constitute unreliable phenotypes. This conclusion corroborates past work on low inter-rater reliability in ICD-10 coding, which may be driven by many factors including patient-provider communication, administrative decision chains, and insurance incentives [8]. Furthermore, these results are consistent with prior work highlighting challenges with UK Biobank clinical codes [20]. Specifically, Stroganov et al highlighted mapping issues of available hospital inpatient and general practitioner information onto the ICD-10 ontology [20]. The limited classification accuracy for most ICD-10 diagnostic groups observed in this study furthermore confirms previous work suggesting that the identification of biomarkers of psychopathology is not feasible without increased efforts to address suboptimal phenotypic reliability [11]. Indeed, a recent study empirically showed the degree to which accuracy of classification models is attenuated as a function of the reliability of the classification target [12]. Notably, age is a phenotype with high reliability and was therefore chosen as a classification target to benchmark our analyses. In summary, our findings reveal limits of the ICD-10 diagnostic ontology that arise from a variety of complex sources, including clinical heterogeneity, lack of inter-rater reliability, and inequities in the structure of healthcare incentives.

Finally, we would like to discuss some methodological limitations of the current study. First, ICD-10 diagnostic groups in the UK Biobank were automatically mapped from available hospital inpatient and general practitioner data sources. These automated mappings are known to be imprecise [20], suggesting that the reliability of ICD-10 diagnostic information in the UK Biobank may be lower than in other research and clinical settings. Second, although we tested our classifications across a range of feature sets and classifiers, it is possible that higher classification accuracy can be achieved using alternative feature sets or imaging modalities (e.g., clinically sensitive modalities such as susceptibility weighted imaging) and/or by using more sensitive classifier architectures (e.g., leveraging deep learning). Third, we identified target diagnostic groups at different levels of the ICD-10 diagnostic hierarchy depending on sample size availability. It is possible that this choice may negatively impact classification accuracy - e.g., by including varying degrees of heterogeneity - although we note that the two diagnostic groups with significant classification results covered both levels of the hierarchy.

### Potential implications

In summary, demyelinating diseases and depression could be classified above chance. These findings were likely driven by a larger magnitude of effect for demyelinating diseases and a larger sample size for depression. Notably, depression was significantly classified based on several functional neuroimaging feature sets but not based on structural feature sets, suggesting that functional neuroimaging features may be more sensitive to depression than structural neuroimaging features. We were unable to reliably classify 15 out of 17 ICD-10 diagnostic cases from matched controls. Overall, these results reveal limits of the ICD-10 ontology. We recommend caution with the use of ICD-10 diagnostic codes as target labels in future work exploring brain-based diagnostic prediction, particularly in epidemiological cohorts like the UK Biobank.

## Methods

### 1. Neuroimaging data and preprocessing

Out of the available UKB neuroimaging data, this work used the T1-weighted scan (1mm isotropic voxels, TR=2000ms, TI = 880ms) and the resting state functional scan (2.4mm isotropic voxels, TR=735ms, TE=39ms, multiband factor 8). Processed data released through the UKB were used (up until and including ICA-FIX clean-up for resting state). T1 and resting state data were transformed into MNI space using the linear and non-linear transforms provided. For detailed preprocessing information, please see [21].

### 2. Classification model

The primary analyses were performed using the Random Forest Classifier as implemented in scikit-learn, informed by prior work [13]. The Random Forest Classifier was selected due to its flexibility in handling data of varied units, its suitability for non-linear classification tasks, and its scalability [22]. Notably, the findings presented in this paper generalized across other classification algorithms (see Methods section 10). Using the pipeline option in scikit-learn, our estimator included scaling, feature space dimension reduction using principal component analysis (PCA), and classification. Nested 5-fold cross-validation was used to tune the PCA dimensionality and the following sets of classifier-specific hyperparameters (Table S3). The depth of the trees and the number of variables considered for splitting were tuned. The number of trees was fixed at 250 following prior work [13, 23]. A shuffle-split resampling scheme was used to subdivide the data into 100 stratified training (80%) and validation (20%) splits. Split validation performance was used to generate the swarm plots.

### 3. Structural neuroimaging feature extraction

Two structural neuroimaging features sets from T1-weighted images were defined based on available imaging derived phenotypes (IDPs) provided by the UKB pipelines [21] (Table S4). The surface-based structural feature set included 285 IDPs available in UKB variable IDs 190 and 196 derived from Freesurfer pipelines. Here, category 196 consists of 186 cortical IDPs from Freesurfer’s DKT-based parcellation and category 190 contains 99 subcortical IDPs (ASEG). The volumetric structural feature set consisted of 153 IDPs from variable IDs 1101 and 1102. Here, category 1101 contains 139 regional gray matter volumes segmented using FSL FAST, and category 1102 contains 14 subcortical volumes segmented using FSL FIRST.

### 4. Alternative feature sets

As a result of the limited classification accuracy based on the primary structural neuroimaging feature sets, we broadened our scope to determine whether alternative feature sets may outperform the primary results. Specifically, we tested 17 different feature sets derived from resting state functional MRI data (Table S4), and one feature set comprised of demographic information (i.e., non-brain data; Table S5). The 17 different resting state functional MRI feature sets reflected combinations across three different brain parcellation (Schaefer parcellation [24], data-driven decomposition using independent component analysis [25, 26], or data-driven decomposition using probabilistic function modes [27, 28]), and across feature types (partial correlation matrix, full correlation matrix, or amplitude [29]), and across dimensionality (ICA only, considering data-driven decomposition dimensionalities of 25, 100, 250, and 300). More detail regarding the resting state functional MRI feature extraction can be found in section 3 of the supplementary materials. The sociodemographic feature set was included based on previous work that revealed that this feature set outperformed neuroimaging-derived features in phenotype prediction [13]. Further details on the sociodemographic feature set can be found in Table S5 of the supplementary materials.

### 5. Statistical Analysis

We used statistical significance as our measure of successful classification. In this study, we computed statistical significance from the distribution of split-wise accuracy scores as the empirical probability of classifying above chance. For a given ICD-10 diagnostic group, a feature set classified significantly above chance if its fitted Student’s *t*-distribution lies, within significance threshold, above the guess line:

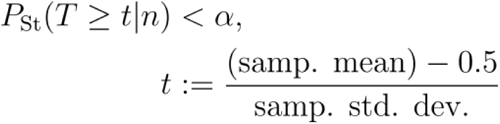

Above, *n* is the number of degrees of freedom (given by the number of shuffle-splits), and we use a significance threshold of *α*=0.05. Note that this computation treats the classification accuracy score of a given shuffle-split as a mean of independent and identically distributed Bernoulli variables and assumes it is asymptotically normally distributed.

We emphasize, however, that we used the sample standard deviation instead of the sample standard error, which makes our significance criterion more stringent than a one-sample *t*-test against 0.5 (chance). A one-sample *t*-test (against chance) would be inappropriate in this context, since it would reflect the likelihood of the population mean accuracy lying above chance, rather than the likelihood of a particular (sub-)population being well-classified in a study on a single dataset. Correction for multiple comparisons was performed across feature sets using the false discovery rate.

### 6. Comparison against multiclass diagnostic classification

In addition to the separate diagnosis-specific case-control classifications, we performed a multiclass classification. For our multiclass classification task, we aimed to categorize samples into 17 ICD-10 groups and the control group (i.e., total of 18 possible labels; see Table S4). Internally, the multiclass procedure trains one classifier for each class, treating the samples of that group as positive and all other samples as negative. The output from the multiclass classifier is combined across all groups. In this multiclass classification, we utilized the unique and matched sample size subject list, as detailed in the case sample selection section to avoid the presence of multiple diagnostic labels per individual case. We employed a Random Forest Classifier with the number of trees set to 250, the criterion for splitting set to “gini”, and the random state set to 42 to ensure reproducibility.

### 7. Comparison against alternative classification models

In addition to the Random Forest Classifier, two further classifiers were tested for classification, namely the Support Vector Classifier and K-Nearest Neighbors Classifier as implemented in scikit-learn. For the Support Vector Classifier, the regularization parameter C and the kernel type were tuned. For the K-Nearest Neighbors Classifier, the number of neighbors, the weight function, and the distance metric were tuned. See Table S3 for hyperparameter values included in the tuning. The classification pipeline described above - including principal component analysis, nested folds for hyperparameter estimation, and shuffle splits - was identical for Random Forest, Support Vector, and K-Nearest Neighbors classifiers.

### 8. Comparison against alternative features sets

In addition to the diagnostic classifications using structural neuroimaging features, we repeated the classification analyses using functional neuroimaging features and socioeconomic features instead. A wide range of options exists to calculate features from resting state functional MRI data [30, 31]. We compared classifications based on feature sets obtained from data-driven approaches including independent component analysis [25, 26] and probabilistic functional modes [27, 28], and atlas-based features [24]. Please see the supplementary methods and Table S4 for further information.

For the socio-economic feature set, we based our selection of features on prior work from [13]. Existing UK Biobank variables in 36 variable IDs across categories of age, sex, education, early life, and lifestyle were selected (see Table S5). Compared to prior work (see Appendix 2, Table S7 in [13]), all variables in the mood & sentiment category and any variables related to smoking behaviors were excluded due to overlap with symptoms commonly observed for several ICD-10 diagnostic groups.

### 9. Comparison against age classification

To benchmark our classification analyses, we repeated the same random forest regression model to classify older versus younger groups based on the same feature sets. To this end, we combined all the subjects from 17 ICD-10 diagnostic groups and subjects with complete neuroimaging data but with no ICD-10 labels in either Chapter V or VI as the cohort. We selected the subjects from this cohort by pairing those with the largest age differences, while ensuring that older and younger groups were matched for sex and head motion. The older subjects were considered as our ‘case’ group (aged 67-70), and the younger subjects were considered the ‘control’ group (aged 40-42). We compiled a balanced group (half “young” and half “old”) of N=2,656 subjects to match the size of the largest ICD-10 diagnostic group, and subsampled N=252 a balanced sublist to match the smallest ICD-10 diagnostic group. This allowed us to benchmark classification effect size across all ICD-10 diagnostic groups against classification effect size of age. To assess the classification accuracy, we employed the same Random Forest classification models (mentioned in the classification model section) on both the structural feature sets (surface and volume) and functional data extracted through Independent Component Analysis, Probabilistic Functional Modes, and the Schaefer atlas (see supplements for details).

## Availability of source code

Project name: WAPIAW3

Project home page: https://github.com/tyo8/WAPIAW3

Operating system: Platform independent

Programming language: Python, shell

License: MIT

## Data availability

UK Biobank data [2, 3] are available following an access application process, for more information please see: https://www.ukbiobank.ac.uk/enable-your-research/apply-for-access. This research was performed under UK Biobank application number 47267.

## Competing interests

The authors declare that they have no competing interests.

## Funding

Janine Bijsterbosch is supported by the NIH (NIMH R01 MH128286 & NIMH R01 MH132962), and Ty Easley is supported by the National Science Foundation under Grant No. DGE-2139839.

## Authors’ contributions

1. Conceptualization: Ty Easley, Janine Bijsterbosch
2. Data Curation: Ty Easley, Petra Lenzini, Xiaoke Luo
3. Formal Analysis: Ty Easley, Xiaoke Luo
4. Funding Acquisition: Janine Bijsterbosch, Ty Easley
5. Methodology: Ty Easley, Xiaoke Luo, Kayla Hannon, Petra Lenzini, Janine Bijsterbosch
6. Supervision: Janine Bijsterbosch
7. Visualization: Xiaoke Luo, Petra Lenzini, Kayla Hannon
8. Writing – Original Draft: Janine Bijsterbosch, Ty Easley, Kayla Hannon, Petra Lenzini, Xiaoke Luo
9. Writing – Review & Editing: Ty Easley, Janine Bijsterbosch, Xiaoke Luo

## Acknowledgments

We are grateful to UK Biobank and the UK Biobank participants for making the resource data possible, and to the data processing team at Oxford University for producing the shared processed data. Thanks to Hailey Modi for pointing us to the scikit-learn pipeline option.

## Notes

### Competing Interest Statement

The authors have declared no competing interest.

