## supplement tables and figures for "Opaque Ontology: Neuroimaging Classification of ICD-10 Diagnostic Groups in the UK Biobank"

Supplementary Materials for ‘Opaque Ontology: Neuroimaging classification of ICD-10 Diagnostic Groups in the UK Biobank’

#### 1. Sample demographics

**Table S1:** Demographics for the surface-based structural analysis diagnostic groups. The age and head motion columns show mean (standard deviation). The sex column reports percent male. Years since diagnosis was obtained by subtracting the date of first occurrence (using UKB variable 41280) from the date of scanning (using UKB variable 53 for instance 2), and dividing the difference in days by 365 (i.e., negative values indicate diagnosis occurred after scanning). Case-control groups were perfectly matched groups for sex (𝞆2 p=1 for all diagnostic groups) and there were no significant group differences for age (p>0.3) nor for head motion (p>0.7). Supplementary Table S2 reveals the same table for unique groups.

| Diagnostic group name | ICD-10 code(s) | N | | Age | | Sex | | Head motion | |
| --- | --- | --- | --- | --- | --- | --- | --- | --- | --- |
|  |  | Case | Control | Case | Control | Case | Control | Case | Control |
| Organic, including symptomatic, mental disorders | F00-F09 | 160 | 160 | 61.44(6.34) | 61.44(6.34) | 61.88% | 61.88% | 0.1568(0.0811) | 0.1566(0.0805) |
| Mental and behavioral disorders due to use of alcohol | F10 | 336 | 336 | 55.34(7.74) | 55.34(7.74) | 75.00% | 75.00% | 0.1482(0.0730) | 0.1477(0.0709) |
| Mental and behavioral disorders due to use of tobacco | F17 | 823 | 823 | 54.40(7.53) | 54.40(7.53) | 56.50% | 56.50% | 0.1487(0.0711) | 0.1478(0.0680) |
| Depressive episode | F32 | 1329 | 1329 | 53.90(7.67) | 53.90(7.67) | 35.36% | 35.36% | 0.1406(0.0695) | 0.1398(0.0665) |
| Other anxiety disorders | F41 | 1045 | 1045 | 54.42(7.55) | 54.42(7.55) | 34.74% | 34.74% | 0.1358(0.0638) | 0.1352(0.0617) |
| Extrapyramidal and movement disorders | G20-G26 | 192 | 192 | 58.77(6.72) | 58.77(6.72) | 52.08% | 52.08% | 0.1382(0.0651) | 0.1381(0.0647) |
| Demyelinating diseases of the central nervous systems | G35-G37 | 125 | 125 | 52.16(7.10) | 52.16(7.10) | 25.60% | 25.60% | 0.1198(0.0592) | 0.1190(0.0575) |
| Epilepsy | G40 | 239 | 239 | 55.62(7.62) | 55.62(7.62) | 47.70% | 47.70% | 0.1394(0.0851) | 0.1376(0.0752) |
| Migraine | G43 | 521 | 521 | 53.46(7.33) | 53.46(7.33) | 25.14% | 25.14% | 0.1257(0.0617) | 0.1247(0.0583) |
| Transient cerebral ischaemic attacks and related syndromes (stroke/TIA) | G45 | 276 | 276 | 59.36(6.03) | 59.36(6.03) | 53.62% | 53.62% | 0.1318(0.0580) | 0.1311(0.0548) |
| Sleep disorders | G47 | 597 | 597 | 55.12(7.18) | 55.12(7.18) | 70.18% | 70.18% | 0.1665(0.0820) | 0.1654(0.0796) |
| Nerve root and plexus compressions in diseases classified elsewhere | G55 | 482 | 482 | 55.39(7.72) | 55.39(7.72) | 49.59% | 49.59% | 0.1352(0.0635) | 0.1349(0.0616) |
| Mononeuropathies of upper limb | G56 | 981 | 981 | 56.26(7.22) | 56.26(7.22) | 32.52% | 32.52% | 0.1414(0.0681) | 0.1407(0.0665) |
| Mononeuropathies of lower limb | G57 | 193 | 193 | 55.13(6.86) | 55.13(6.86) | 21.76% | 21.76% | 0.1246(0.0608) | 0.1240(0.0582) |
| Other polyneuropathies | G62 | 157 | 157 | 58.38(6.70) | 58.38(6.70) | 54.78% | 54.78% | 0.1412(0.0710) | 0.1390(0.0624) |
| Cerebral Palsy and other paralytic syndromes | G80-G83 | 158 | 158 | 56.83(8.12) | 56.83(8.12) | 55.06% | 55.06% | 0.1273(0.0555) | 0.1271(0.0554) |
| Other disorders of brain | G93 | 148 | 148 | 54.52(8.20) | 54.52(8.20) | 39.86% | 39.86% | 0.1317(0.0530) | 0.1307(0.0509) |

**Table S2:** Demographics for the unique diagnostic groups. Diagnostic group demographics. The age and head motion columns show mean (± standard deviation). The sex column reports percent recorded as male. * indicates a significant (p<0.05) case-control difference within the diagnostic group.

| Diagnostic group name | ICD-10 code | N | Age | Sex | Head motion |
| --- | --- | --- | --- | --- | --- |
| Organic, including symptomatic, mental disorders | F00-F09 | 59 | 61.39(6.25) | 55.93% | 0.1524(0.0677) |
| Mental and behavioral disorders due to use of alcohol | F10 | 59 | 57.10(7.26) | 33.90% | 0.1223(0.0505) |
| Mental and behavioral disorders due to use of tobacco | F17 | 59 | 56.71(7.85) | 33.90% | 0.1282(0.0436) |
| Depressive episode | F32 | 59 | 56.73(7.88) | 33.90% | 0.1230(0.0412) |
| Other anxiety disorders | F41 | 59 | 56.73(7.88) | 33.90% | 0.1230(0.0409) |
| Extrapyramidal and movement disorders | G20-G26 | 59 | 57.98(6.47) | 33.90% | 0.1135(0.0385) |
| Demyelinating diseases of the central nervous systems | G35-G37 | 59 | 54.05(6.98) | 33.90% | 0.1305(0.0675) |
| Epilepsy | G40 | 59 | 56.68(7.83) | 33.90% | 0.1359(0.0602) |
| Migraine | G43 | 59 | 56.69(7.85) | 33.90% | 0.1169(0.0372) |
| Transient cerebral ischaemic attacks and related syndromes (stroke/TIA) | G45 | 59 | 57.36(7.06) | 33.90% | 0.1213(0.0547) |
| Sleep disorders | G47 | 59 | 56.73(7.73) | 33.90% | 0.1360(0.0501) |
| Nerve root and plexus compressions in diseases classified elsewhere | G55 | 59 | 56.71(7.88) | 33.90% | 0.1172(0.0336) |
| Mononeuropathies of upper limb | G56 | 59 | 56.73(7.88) | 33.90% | 0.1345(0.0506) |
| Mononeuropathies of lower limb | G57 | 59 | 56.27(7.39) | 33.90% | 0.1253(0.0530) |
| Other polyneuropathies | G62 | 59 | 58.98(6.75) | 33.90% | 0.1251(0.0581) |
| Cerebral Palsy and other paralytic syndromes | G80-G83 | 59 | 56.92(7.69) | 47.46% | 0.1203(0.0473) |
| Other disorders of brain | G93 | 59 | 56.73(7.88) | 33.90% | 0.1263(0.0447) |
| Health | - | 59 | 56.75(7.84) | 33.90% | 0.1259(0.0418) |

#### 2. Classification model hyperparameters

**Table S3:** Hyperparameter search grids for all scikit-learn classification models. Note that “rank” hear means “maximum possible rank” and is given by min(*n*,*m*) for
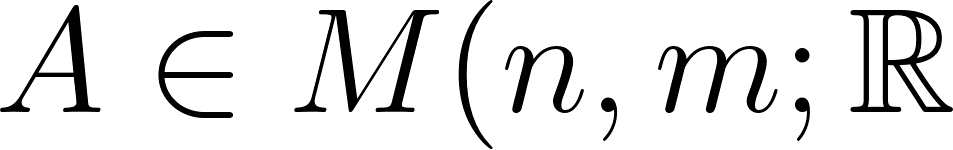
); rank was not estimated for each input dataset.

| **Random Forest Classification** | |
| --- | --- |
| **Hyperparameter** | **Values** |
| Number of PCs | rank/100, rank/32, rank/10, rank/3.2, full rank |
| Impurity criterion | Gini |
| Maximum tree depth | 5, 10, 20, 40, full depth |
| Fraction of features for split | 1, 5, log2, sqrt, complete |
| Number of trees | 250 (fixed) |
| **Support Vector Classification** | |
| **Hyperparameter** | **Values** |
| Number of PCs | rank/100, rank/32, rank/10, rank/3.2, full rank |
| Margin regularization coefficient *C* | 10^-3^, 10^-2^, 10^-1^, 1, 10^1^, 10^2^, 10^3^ |
| Kernel function | ‘linear’, ‘rbf’, ‘sigmoid’, ‘poly’ |
| **K-Nearest Neighbor Classification** | |
| **Hyperparameter** | **Values** |
| Number of PCs | rank/100, rank/32, rank/10, rank/3.2, full rank |
| Number of neighbors | 1, 5, 11, 18, 27 |
| Neighbor weights | ‘uniform’, ‘distance’ |
| Distance metric | ‘L1’, ‘L2’, ‘cosine’ |

#### 3. Resting state feature extraction

Resting state feature extraction was performed specifically for this work as described in section 4(a-d) below. As such, none of the UKB resting state imaging derived phenotypes (IDPs) were used. The reasons for not using the existing independent component analysis (ICA) IDPs were twofold. First, we did not want the network estimation to be influenced (and potentially biased) by participants not included in this study. Second, because we set out to test additional brain rfMRI representation choices, the UKB’s precomputed resting-state IDPs did not provide sufficient variety or coverage. Structural features were selected from the existing UKB IDPs because these are estimated at the single-subject level (without group input and therefore without potential bias; see section 4e for further details). An overview of all feature sets is shown in Table S3.

**Table S4:** Overview of 2 structural MRI feature sets, 16 resting state functional MRI feature sets, and 1 sociodemographic feature set. Parcellation dimensionality is in regions/network within the whole-brain map unless otherwise stated.

| **Brain representation** | **Feature set** | **Parcellation dimensionality** | **N features** |
| --- | --- | --- | --- |
| Structural | Surface  (= 1 feature set) | Freesurfer DKT (186)  Freesurfer ASEG (190) | 285 |
|  | Volume  (= 1 feature set) | FAST regional GMV (139)  FIRST subcortical GMV (14) | 153 |
| Schaefer | Correlation matrix (partial or full)  (= 2 features sets | 400 | 79,800 |
|  | Amplitudes  (= 1 feature set) | 400 | 400 |
| ICA | Correlation matrix (partial or full)  (= 8 feature sets) | 25 | 300 |
|  |  | 100 | 4,950 |
|  |  | 150 | 11,175 |
|  |  | 300 | 44,850 |
|  | Amplitudes  (=4 feature sets) | 25 | 25 |
|  |  | 100 | 100 |
|  |  | 150 | 150 |
|  |  | 300 | 300 |
| PFM | Spatial overlap matrix,  Correlation matrix (full)  (=2 feature sets) | 25 | 300 |
| Sociodemographics | See Table S5  (= 1 feature set) | N.A. | 36 |

##### a. Independent component analysis (ICA)

Group ICA was performed separately within each of the 17 diagnostic groups, using the combined cases and controls subject list (and also repeated separately for the unique diagnostic groups in which overlapping cases and their matched controls are removed). ICA dimensionalities of 25, 100, 150 and 300 were performed. The first two dimensionalities (25 and 100) were chosen to match the dimensionality of released IDPs. Following dual regression to obtain participant-specific timeseries for each network, full and partial correlation features were each used for classification. Following dual regression, the standard deviations of participant-specific network timeseries (i.e., amplitudes) were used for classification.

##### b. Probabilistic Functional Modes (PFM)

Probabilistic Functional Modes (PFM) is a hierarchical Bayesian algorithm that was developed as an alternative to the group ICA and dual regression pipeline described above (1). Advantages of PFM relative to ICA include the hierarchical framework that optimizes group and participant solutions iteratively to improve sensitivity to individual differences (2), and the relaxation of the spatial independence constraint to potentially allow for greater network overlap (3). As with the ICA pipeline above, PFM was performed separately within each of the 17 diagnostic groups, using the combined cases and controls subject list. PFM was performed at a dimensionality of 25 to match the lowest ICA dimension. PFM was not performed at higher dimensions due to the high demand on computing resources (4). Using participant-specific spatial maps, the spatial overlap matrix features (full correlation across vectorized spatial maps) were used for classification (3).

##### c. Schaefer atlas

Timeseries were extracted using the 400 dimensionality Schaefer atlas (5). Since timeseries extraction is performed independently for each participant, no separate feature extraction per diagnostic group was required. Per-subject partial and full correlation features (upper triangle of the 400x400 matrix) were each used for classification.

#### 4. Sociodemographic features

**Table S5:** Information regarding socio demographic features. The 36 sociodemographic variables listed below constitute a 36-dimensional predictive feature set after one-hot encoding categorical variables.

| **Category** | **UKB variable ID** | **Description** |
| --- | --- | --- |
| Age, sex | 31-0.0  34-0.0  52-0.0  21022-0.0  21003-2.0 | Sex  Year of birth  Month of birth  Age at recruitment  Age when attended assessment centre |
| Education | 6138-2.0  845-2.0 | Qualifications  Age completed full time education |
| Early life | 1647-2.0  1677-2.0 1687-2.0  1697-2.0 1707-2.0  1767-2.0  1777-2.0  1787-2.0 | Country of birth (UK/elsewhere)  Breastfed as a baby  Comparative body size at age 10  Comparative height size at age 10  Handedness (chirality/laterality)  Adopted as a child  Part of a multiple birth  Maternal smoking around birth |
| Lifestyle | 670-2.0  680-2.0  6139-2.0  699-2.0  709-2.0  6141-2.0  728-2.0  738-2.0  796-2.0  757-2.0  767-2.0  777-2.0  6143-2.0  6142-2.0  806-2.0  816-2.0  826-2.0  3426-2.0  1031-2.0  6160-2.0  2110-2.0 | Type of accommodation lived in  Own or rent accommodation lived in  Gas or solid-fuel cooking/heating  Length of time at current address  Number in household  How are people in household related to participant  Number of vehicles in household  Income before tax  Distance between home and job workplace  Time employed in main current job  Length of working week for main job  Freq. of traveling from home to job workplace Transport type for commuting to job workplace  Current employment status  Job involves mainly walking or standing  Job involves heavy manual or physical work  Job involves shift work  Job involves night shift work  Freq. of friend/ family visits  Leisure/social activities  Able to confide |

### Supplementary Results

**Figure S1:** Volumetric structural results

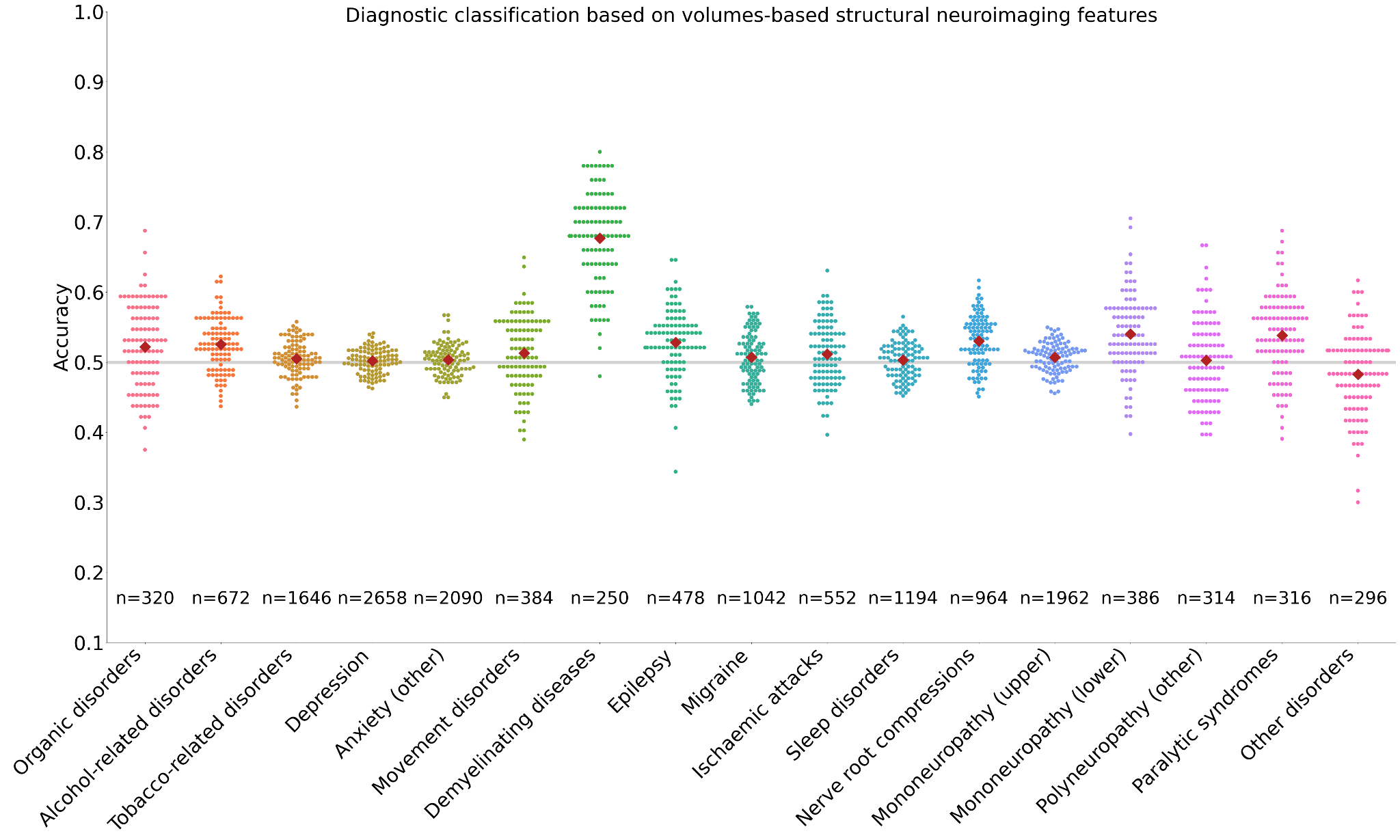

**Figure S2:** Multiclass volumetric results

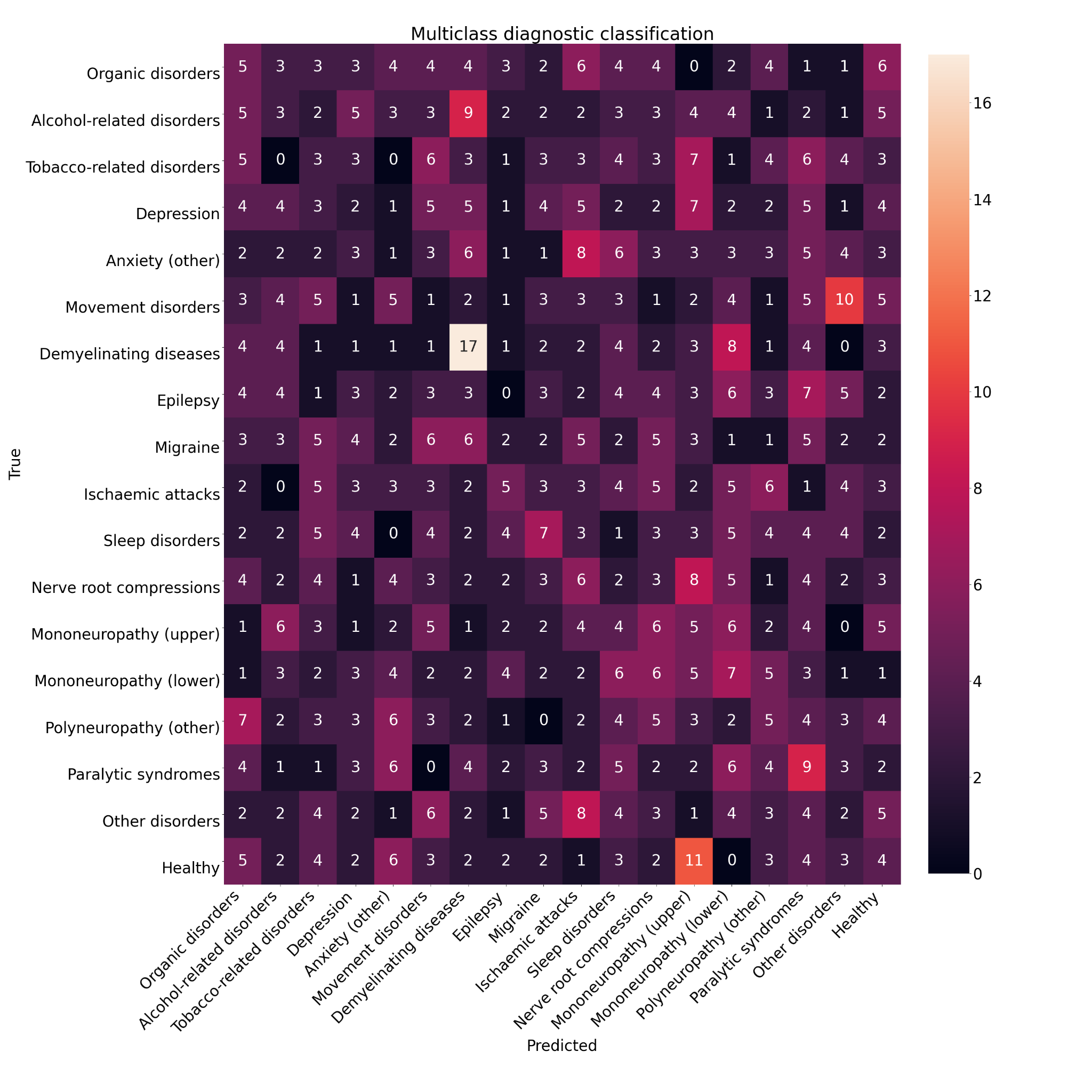

**Table S6:** Overview of primary Random Forest Classification results from structural neuroimaging data only. Each cell reports to mean accuracy (Ac), uncorrected p-value (pu), and corrected p-value (pc). Corrected p-values were corrected for multiple comparisons across 2 feature sets (i.e., within each column) using false discovery rate correction. Cells highlighted in green indicate significant results (pc< 0.05) and cells highlighted in yellow indicate trend-level results (0.05 < pC < 0.10).

|  | | ***F00-***  ***F09*** | ***F10*** | ***F17*** | ***F32*** | ***F41*** | ***G20-***  ***G26*** | ***G35-***  ***G37*** | ***G40*** | ***G43*** | ***G45*** | ***G47*** | ***G55*** | ***G56*** | ***G57*** | ***G62*** | ***G80-***  ***G83*** | ***G93*** |
| --- | --- | --- | --- | --- | --- | --- | --- | --- | --- | --- | --- | --- | --- | --- | --- | --- | --- | --- |
| **RFC** | ***Surf.*** | **Ac**=.55  ***p_u_***=0.17  ***p_c_***=0.70 | **Ac**=.54  ***p_u_***=0.13  ***p_c_***=0.59 | **Ac**=.51  ***p_u_***=0.35  ***p_c_***=0.33 | **Ac**=.52  ***p_u_***=0.19  ***p_c_***=0.24 | **Ac**=.51  ***p_u_***=0.31  ***p_c_***=0.42 | **Ac**=.53  ***p_u_***=0.30  ***p_c_***=0.67 | **Ac**=.63  ***p_u_***=0.01  ***p_c_***=0.13 | **Ac**=.50  ***p_u_***=0.48  ***p_c_***=0.52 | **Ac**=.51  ***p_u_***=0.39  ***p_c_***=0.58 | **Ac**=.50  ***p_u_***=0.52  ***p_c_***=0.66 | **Ac**=.51  ***p_u_***=0.41  ***p_c_***=0.48 | **Ac**=.51  ***p_u_***=0.40  ***p_c_***=0.55 | **Ac**=.51  ***p_u_***=0.32  ***p_c_***=0.52 | **Ac**=.50  ***p_u_***=0.49  ***p_c_***=0.55 | **Ac**=.52  ***p_u_***=0.34  ***p_c_***=0.67 | **Ac**=.49  ***p_u_***=0.55  ***p_c_***=0.61 | **Ac**=.48  ***p_u_***=0.61  ***p_c_***=0.64 |
|  | ***Vol.*** | **Ac**=.52  ***p_u_***=0.36  ***p_c_***=0.70 | **Ac**=.53  ***p_u_***=0.25  ***p_c_***=0.59 | **Ac**=.51  ***p_u_***=0.42  ***p_c_***=0.33 | **Ac**=.50  ***p_u_***=0.46  ***p_c_***=0.48 | **Ac**=.50  ***p_u_***=0.44  ***p_c_***=0.52 | **Ac**=.51  ***p_u_***=0.40  ***p_c_***=0.67 | **Ac**=.68  ***p_u_***=4e-3  ***p_c_***=0.09 | **Ac**=.53  ***p_u_***=0.28  ***p_c_***=0.51 | **Ac**=.51  ***p_u_***=0.42  ***p_c_***=0.58 | **Ac**=.51  ***p_u_***=0.40  ***p_c_***=0.66 | **Ac**=.50  ***p_u_***=0.45  ***p_c_***=0.48 | **Ac**=.53  ***p_u_***=0.20  ***p_c_***=0.55 | **Ac**=.51  ***p_u_***=0.37  ***p_c_***=0.52 | **Ac**=.54  ***p_u_***=0.24  ***p_c_***=0.55 | **Ac**=.50  ***p_u_***=0.48  ***p_c_***=0.67 | **Ac**=.54  ***p_u_***=0.26  ***p_c_***=0.61 | **Ac**=.48  ***p_u_***=0.61  ***p_c_***=0.64 |

**Table S7:** Overview of primary Random Forest Classification results from structural neuroimaging data using the matched sample sizes (i.e., 125 cases and 125 controls in each ICD-10 diagnostic category). Each cell reports to mean accuracy (Ac), uncorrected p-value (pu), and corrected p-value (pc). Corrected p-values were corrected for multiple comparisons across 2 feature sets (i.e., within each column) using false discovery rate correction. Cells highlighted in green indicate significant results (pc< 0.05) and cells highlighted in yellow indicate trend-level results (0.05 < pC < 0.10).

|  | | ***F00-***  ***F09*** | ***F10*** | ***F17*** | ***F32*** | ***F41*** | ***G20-***  ***G26*** | ***G35-***  ***G37*** | ***G40*** | ***G43*** | ***G45*** | ***G47*** | ***G55*** | ***G56*** | ***G57*** | ***G62*** | ***G80-***  ***G83*** | ***G93*** |
| --- | --- | --- | --- | --- | --- | --- | --- | --- | --- | --- | --- | --- | --- | --- | --- | --- | --- | --- |
| **RFC** | ***Surf.*** | **Ac**=.56  ***p_u_***=0.18  ***p_c_***=0.18 | **Ac**=.53  ***p_u_***=0.31  ***p_c_***=0.31 | **Ac**=.54  ***p_u_***=0.24  ***p_c_***=0.24 | **Ac**=.53  ***p_u_***=0.31  ***p_c_***=0.31 | **Ac**=.49  ***p_u_***=0.57  ***p_c_***=0.57 | **Ac**=.50  ***p_u_***=0.53  ***p_c_***=0.53 | **Ac**=.64  ***p_u_***=0.01  ***p_c_***=0.01 | **Ac**=.50  ***p_u_***=0.49  ***p_c_***=0.49 | **Ac**=.49  ***p_u_***=0.58  ***p_c_***=0.58 | **Ac**=.50  ***p_u_***=0.53  ***p_c_***=0.53 | **Ac**=.50  ***p_u_***=0.52  ***p_c_***=0.52 | **Ac**=.52  ***p_u_***=0.36  ***p_c_***=0.36 | **Ac**=.49  ***p_u_***=0.55  ***p_c_***=0.55 | **Ac**=.50  ***p_u_***=0.53  ***p_c_***=0.53 | **Ac**=.50  ***p_u_***=0.49  ***p_c_***=0.49 | **Ac**=.50  ***p_u_***=0.49  ***p_c_***=0.49 | **Ac**=.48  ***p_u_***=0.64  ***p_c_***=0.64 |
|  | ***Vol.*** | **Ac**=.53  ***p_u_***=0.32  ***p_c_***=0.32 | **Ac**=.51  ***p_u_***=0.42  ***p_c_***=0.42 | **Ac**=.52  ***p_u_***=0.39  ***p_c_***=0.39 | **Ac**=.49  ***p_u_***=0.57  ***p_c_***=0.57 | **Ac**=.50  ***p_u_***=0.50  ***p_c_***=0.50 | **Ac**=.51  ***p_u_***=0.47  ***p_c_***=0.47 | **Ac**=.69  ***p_u_***=2e-3  ***p_c_***=2e-3 | **Ac**=.53  ***p_u_***=0.29  ***p_c_***=0.29 | **Ac**=.48  ***p_u_***=0.60  ***p_c_***=0.60 | **Ac**=.50  ***p_u_***=0.48  ***p_c_***=0.48 | **Ac**=.54  ***p_u_***=0.27  ***p_c_***=0.27 | **Ac**=.51  ***p_u_***=0.45  ***p_c_***=0.45 | **Ac**=.52  ***p_u_***=0.36  ***p_c_***=0.36 | **Ac**=.51  ***p_u_***=0.45  ***p_c_***=0.45 | **Ac**=.49  ***p_u_***=0.54  ***p_c_***=0.54 | **Ac**=.51  ***p_u_***=0.47  ***p_c_***=0.47 | **Ac**=.48  ***p_u_***=0.64  ***p_c_***=0.64 |

**Table S8**: Overview of results over varying classifiers. Support Vector (SVC) and k-Nearest Neighbors Classification (KNC) of diagnostic codes from structural neuroimaging features. In this experiment, we corrected for a total of 6 multiple comparisons over classifier choice and structural feature type. The results do not change: only G35-37 (Demyelinating diseases; Table S1) is classified significantly above chance from structural features.

|  | | ***F00-***  ***F09*** | ***F10*** | ***F17*** | ***F32*** | ***F41*** | ***G20-***  ***G26*** | ***G35-***  ***G37*** | ***G40*** | ***G43*** | ***G45*** | ***G47*** | ***G55*** | ***G56*** | ***G57*** | ***G62*** | ***G80-***  ***G83*** | ***G93*** |
| --- | --- | --- | --- | --- | --- | --- | --- | --- | --- | --- | --- | --- | --- | --- | --- | --- | --- | --- |
| **RFC** | ***Surf.*** | **Ac**=.55  ***p_u_***=0.17  ***p_c_***=0.46 | **Ac**=.54  ***p_u_***=0.13  ***p_c_***=0.27 | **Ac**=.51  ***p_u_***=0.35  ***p_c_***=0.42 | **Ac**=.52  ***p_u_***=0.19  ***p_c_***=0.60 | **Ac**=.51  ***p_u_***=0.31  ***p_c_***=0.51 | **Ac**=.53  ***p_u_***=0.30  ***p_c_***=0.40 | **Ac**=.63  ***p_u_***=0.01  ***p_c_***=0.09 | **Ac**=.50  ***p_u_***=0.48  ***p_c_***=0.55 | **Ac**=.51  ***p_u_***=0.39  ***p_c_***=0.60 | **Ac**=.50  ***p_u_***=0.52  ***p_c_***=0.55 | **Ac**=.51  ***p_u_***=0.41  ***p_c_***=0.55 | **Ac**=.51  ***p_u_***=0.40  ***p_c_***=0.40 | **Ac**=.51  ***p_u_***=0.32  ***p_c_***=0.58 | **Ac**=.50  ***p_u_***=0.49  ***p_c_***=0.54 | **Ac**=.52  ***p_u_***=0.34  ***p_c_***=0.49 | **Ac**=.49  ***p_u_***=0.55  ***p_c_***=0.61 | **Ac**=.48  ***p_u_***=0.61  ***p_c_***=0.61 |
|  | ***Vol.*** | **Ac**=.52  ***p_u_***=0.36  ***p_c_***=0.46 | **Ac**=.53  ***p_u_***=0.25  ***p_c_***=0.27 | **Ac**=.51  ***p_u_***=0.42  ***p_c_***=0.42 | **Ac**=.50  ***p_u_***=0.46  ***p_c_***=0.60 | **Ac**=.50  ***p_u_***=0.44  ***p_c_***=0.51 | **Ac**=.51  ***p_u_***=0.40  ***p_c_***=0.40 | **Ac**=.68  ***p_u_***=4e-3  ***p_c_***=0.02 | **Ac**=.53  ***p_u_***=0.28  ***p_c_***=0.55 | **Ac**=.51  ***p_u_***=0.42  ***p_c_***=0.60 | **Ac**=.51  ***p_u_***=0.40  ***p_c_***=0.55 | **Ac**=.50  ***p_u_***=0.45  ***p_c_***=0.55 | **Ac**=.53  ***p_u_***=0.20  ***p_c_***=0.40 | **Ac**=.51  ***p_u_***=0.37  ***p_c_***=0.58 | **Ac**=.54  ***p_u_***=0.24  ***p_c_***=0.54 | **Ac**=.50  ***p_u_***=0.48  ***p_c_***=0.49 | **Ac**=.54  ***p_u_***=0.26  ***p_c_***=0.61 | **Ac**=.48  ***p_u_***=0.61  ***p_c_***=0.61 |
| **SVC** | ***Surf.*** | **Ac**=0.55  ***p_u_***=0.17  ***p_c_***=0.46 | **Ac**=0.54  ***p_u_***=0.13  ***p_c_***=0.27 | **Ac**=0.52  ***p_u_***=0.23  ***p_c_***=0.42 | **Ac**=0.52  ***p_u_***=0.19  ***p_c_***=0.60 | **Ac**=0.51  ***p_u_***=0.31  ***p_c_***=0.51 | **Ac**=0.53  ***p_u_***=0.28  ***p_c_***=0.40 | **Ac**=0.67  ***p_u_***=4e-3  ***p_c_***=0.02 | **Ac**=0.52  ***p_u_***=0.35  ***p_c_***=0.55 | **Ac**=0.52  ***p_u_***=0.31  ***p_c_***=0.60 | **Ac**=0.50  ***p_u_***=0.47  ***p_c_***=0.55 | **Ac**=0.50  ***p_u_***=0.53  ***p_c_***=0.55 | **Ac**=0.51  ***p_u_***=0.33  ***p_c_***=0.40 | **Ac**=0.50  ***p_u_***=0.58  ***p_c_***=0.58 | **Ac**=0.50  ***p_u_***=0.52  ***p_c_***=0.54 | **Ac**=0.54  ***p_u_***=0.25  ***p_c_***=0.49 | **Ac**=0.50  ***p_u_***=0.48  ***p_c_***=0.61 | **Ac**=0.49  ***p_u_***=0.59  ***p_c_***=0.61 |
|  | ***Vol.*** | **Ac**=0.54  ***p_u_***=0.25  ***p_c_***=0.46 | **Ac**=0.52  ***p_u_***=0.24  ***p_c_***=0.27 | **Ac**=0.53  ***p_u_***=0.17  ***p_c_***=0.42 | **Ac**=0.51  ***p_u_***=0.29  ***p_c_***=0.60 | **Ac**=0.51  ***p_u_***=0.40  ***p_c_***=0.51 | **Ac**=0.52  ***p_u_***=0.33  ***p_c_***=0.40 | **Ac**=0.68  ***p_u_***=1e-3  ***p_c_***=9e-3 | **Ac**=0.54  ***p_u_***=0.20  ***p_c_***=0.55 | **Ac**=0.52  ***p_u_***=0.31  ***p_c_***=0.60 | **Ac**=0.49  ***p_u_***=0.55  ***p_c_***=0.55 | **Ac**=0.51  ***p_u_***=0.34  ***p_c_***=0.55 | **Ac**=0.51  ***p_u_***=0.32  ***p_c_***=0.40 | **Ac**=0.51  ***p_u_***=0.26  ***p_c_***=0.58 | **Ac**=0.53  ***p_u_***=0.26  ***p_c_***=0.54 | **Ac**=0.50  ***p_u_***=0.49  ***p_c_***=0.49 | **Ac**=0.53  ***p_u_***=0.26  ***p_c_***=0.61 | **Ac**=.49  ***p_u_***=0.55  ***p_c_***=0.61 |
| **KNC** | ***Surf.*** | **Ac**=0.53  ***p_u_***=0.28  ***p_c_***=0.46 | **Ac**=0.54  ***p_u_***=0.15  ***p_c_***=0.27 | **Ac**=0.51  ***p_u_***=0.34  ***p_c_***=0.42 | **Ac**=0.50  ***p_u_***=0.59  ***p_c_***=0.60 | **Ac**=0.50  ***p_u_***=0.51  ***p_c_***=0.51 | **Ac**=0.53  ***p_u_***=0.25  ***p_c_***=0.40 | **Ac**=0.59  ***p_u_***=0.11  ***p_c_***=0.11 | **Ac**=0.49  ***p_u_***=0.55  ***p_c_***=0.55 | **Ac**=0.51  ***p_u_***=0.33  ***p_c_***=0.60 | **Ac**=0.51  ***p_u_***=0.38  ***p_c_***=0.55 | **Ac**=0.50  ***p_u_***=0.47  ***p_c_***=0.55 | **Ac**=0.51  ***p_u_***=0.38  ***p_c_***=0.40 | **Ac**=0.50  ***p_u_***=0.48  ***p_c_***=0.58 | **Ac**=0.49  ***p_u_***=0.54  ***p_c_***=0.54 | **Ac**=0.50  ***p_u_***=0.48  ***p_c_***=0.49 | **Ac**=0.48  ***p_u_***=0.61  ***p_c_***=0.61 | **Ac**=0.50  ***p_u_***=0.52  ***p_c_***=0.61 |
|  | ***Vol.*** | **Ac**=0.51  ***p_u_***=0.46  ***p_c_***=0.46 | **Ac**=0.52  ***p_u_***=0.27  ***p_c_***=0.27 | **Ac**=0.52  ***p_u_***=0.27  ***p_c_***=0.42 | **Ac**=0.50  ***p_u_***=0.60  ***p_c_***=0.60 | **Ac**=0.50  ***p_u_***=0.51  ***p_c_***=0.51 | **Ac**=0.52  ***p_u_***=0.35  ***p_c_***=0.40 | **Ac**=0.61  ***p_u_***=0.06  ***p_c_***=0.11 | **Ac**=0.53  ***p_u_***=0.26  ***p_c_***=0.55 | **Ac**=0.49  ***p_u_***=0.60  ***p_c_***=0.60 | **Ac**=0.50  ***p_u_***=0.51  ***p_c_***=0.55 | **Ac**=0.50  ***p_u_***=0.55  ***p_c_***=0.55 | **Ac**=0.52  ***p_u_***=0.26  ***p_c_***=0.40 | **Ac**=0.50  ***p_u_***=0.58  ***p_c_***=0.58 | **Ac**=0.52  ***p_u_***=0.32  ***p_c_***=0.54 | **Ac**=0.50  ***p_u_***=0.49  ***p_c_***=0.49 | **Ac**=0.49  ***p_u_***=0.54  ***p_c_***=0.61 | **Ac**=0.48  ***p_u_***=0.61  ***p_c_***=0.61 |

**Table S9:** Overview of all Random Forest Classification results. Complete table summarizing all random forest classification results. Each cell reports to mean accuracy (Ac), uncorrected p-value (pu), and corrected p-value (pc). Corrected p-values were corrected for multiple comparisons across 20 feature sets (i.e., within each row) using false discovery rate correction. Cells highlighted in green indicate significant results (pc< 0.05) and cells highlighted in yellow indicate trend-level results (0.05 < pC < 0.10). Age prediction was not performed for PROFUMO and sociodemographic results as indicated in the cells highlighted in black. Surf. = surface. Amps = amplitudes. Fnets = full correlation network matrix. Pnets = partial correlation network matrix. ICA = independent component analysis. Schaef. = Schaefer parcellation. PROF. = PROFUMO. SpNets = spatial correlation overlap matrix. Only F32 (Depression; see Table S1) was classified significantly above chance after multiple comparisons correction.

| **ICD-10 code** | **Structural** | | | **Functional** | | | | | | | | | | | | | | | | | ***Socio-***  ***demo-***  ***graphic*** |
| --- | --- | --- | --- | --- | --- | --- | --- | --- | --- | --- | --- | --- | --- | --- | --- | --- | --- | --- | --- | --- | --- |
|  | ***Surf.*** | ***Surf.***  ***(same size)*** | ***Volume*** | ***ICA 25***  ***Amps*** | ***ICA 25***  ***Fnets*** | ***ICA 25***  ***Pnets*** | ***ICA 100***  ***Amps*** | ***ICA 100***  ***Fnets*** | ***ICA 100***  ***Pnets*** | ***ICA 150***  ***Amps*** | ***ICA 150***  ***Fnets*** | ***ICA 150***  ***Pnets*** | ***ICA 300***  ***Amps*** | ***ICA 300***  ***Fnets*** | ***ICA 300***  ***Pnets*** | ***SchaefAmps*** | ***SchaefFnets*** | ***SchaefPnets*** | ***PROF. SpNets*** | ***PROF. Fnets*** |  |
| ***F00-***  ***F09*** | Ac=.55  p_u_=0.17  p_c_=0.70 | Ac=.56  p_u_=0.18  p_c_=0.18 | Ac=.52  p_u_=0.36  p_c_=0.70 | Ac=.48  p_u_=0.63  p_c_=0.70 | Ac=.47  p_u_=0.63  p_c_=0.70 | Ac=.51  p_u_=0.47  p_c_=0.70 | Ac=.50  p_u_=0.50  p_c_=0.70 | Ac=.49  p_u_=0.58  p_c_=0.70 | Ac=.49  p_u_=0.59  p_c_=0.70 | Ac=.50  p_u_=0.52  p_c_=0.70 | Ac=.47  p_u_=0.77  p_c_=0.70 | Ac=.50  p_u_=0.53  p_c_=0.70 | Ac=.48  p_u_=0.63  p_c_=0.70 | Ac=.51  p_u_=0.44  p_c_=0.70 | Ac=.49  p_u_=0.54  p_c_=0.70 | Ac=.50  p_u_=0.47  p_c_=0.70 | Ac=.52  p_u_=0.43  p_c_=0.70 | Ac=.51  p_u_=0.43  p_c_=0.70 | Ac=.51  p_u_=0.45  p_c_=0.70 | Ac=.52  p_u_=0.37  p_c_=0.70 | Ac=.52  p_u_=0.36  p_c_=0.70 |
| ***F10*** | Ac=.54  p_u_=0.13  p_c_=0.59 | Ac=.53  p_u_=0.31  p_c_=0.31 | Ac=.53  p_u_=0.25  p_c_=0.59 | Ac=.51  p_u_=0.44  p_c_=0.59 | Ac=.50  p_u_=0.47  p_c_=0.59 | Ac=.51  p_u_=0.36  p_c_=0.59 | Ac=.51  p_u_=0.40  p_c_=0.59 | Ac=.52  p_u_=0.30  p_c_=0.59 | Ac=.50  p_u_=0.59  p_c_=0.59 | Ac=.52  p_u_=0.34  p_c_=0.59 | Ac=.52  p_u_=0.31  p_c_=0.59 | Ac=.51  p_u_=0.39  p_c_=0.59 | Ac=.50  p_u_=0.46  p_c_=0.59 | Ac=.52  p_u_=0.33  p_c_=0.59 | Ac=.49  p_u_=0.58  p_c_=0.59 | Ac=.50  p_u_=0.48  p_c_=0.59 | Ac=.52  p_u_=0.27  p_c_=0.59 | Ac=.50  p_u_=0.53  p_c_=0.59 | Ac=.50  p_u_=0.50  p_c_=0.59 | Ac=.50  p_u_=0.47  p_c_=0.59 | Ac=.53  p_u_=0.17  p_c_=0.59 |
| ***F17*** | Ac=.51  p_u_=0.35  p_c_=0.33 | Ac=.54  p_u_=0.24  p_c_=0.24 | Ac=.51  p_u_=0.42  p_c_=0.33 | Ac=.52  p_u_=0.17  p_c_=0.37 | Ac=.52  p_u_=0.28  p_c_=0.33 | Ac=.52  p_u_=0.23  p_c_=0.35 | Ac=.52  p_u_=0.26  p_c_=0.33 | Ac=.52  p_u_=0.18  p_c_=0.33 | Ac=.53  p_u_=0.13  p_c_=0.33 | Ac=.54  p_u_=0.12  p_c_=0.37 | Ac=.53  p_u_=0.08  p_c_=0.33 | Ac=.52  p_u_=0.17  p_c_=0.33 | Ac=.52  p_u_=0.20  p_c_=0.33 | Ac=.53  p_u_=0.10  p_c_=0.33 | Ac=.50  p_u_=0.55  p_c_=0.33 | Ac=.52  p_u_=0.19  p_c_=0.33 | Ac=.53  p_u_=0.11  p_c_=0.33 | Ac=.50  p_u_=0.59  p_c_=0.33 | Ac=.53  p_u_=0.16  p_c_=0.33 | Ac=.50  p_u_=0.43  p_c_=0.33 | Ac=.57  p_u_=4e-3  p_c_=0.07 |
| ***F32*** | Ac=.52  p_u_=0.19  p_c_=0.24 | Ac=.53  p_u_=0.31  p_c_=0.31 | Ac=.50  p_u_=0.46  p_c_=0.48 | Ac=.54  p_u_=0.01  p_c_=0.06 | Ac=.54  p_u_=0.05  p_c_=0.10 | Ac=.52  p_u_=0.14  p_c_=0.20 | Ac=.54  p_u_=0.06  p_c_=0.10 | Ac=.54  p_u_=0.02  p_c_=0.06 | Ac=.53  p_u_=0.05  p_c_=0.10 | Ac=.54  p_u_=0.02  p_c_=0.06 | Ac=.55  p_u_=0.01  p_c_=0.05 | Ac=.52  p_u_=0.18  p_c_=0.24 | Ac=.53  p_u_=0.04  p_c_=0.10 | Ac=.54  p_u_=3e-3  p_c_=0.02 | Ac=.50  p_u_=0.43  p_c_=0.48 | Ac=.52  p_u_=0.14  p_c_=0.20 | Ac=.55  p_u_=3e-3  p_c_=0.02 | Ac=.49  p_u_=0.59  p_c_=0.59 | Ac=.53  p_u_=0.06  p_c_=0.10 | Ac=.51  p_u_=0.32  p_c_=0.38 | Ac=.58  p_u_=5e-5  p_c_=1e-3 |
| ***F41*** | Ac=.51  p_u_=0.31  p_c_=0.42 | Ac=.49  p_u_=0.57  p_c_=0.57 | Ac=.50  p_u_=0.44  p_c_=0.52 | Ac=.51  p_u_=0.25  p_c_=0.39 | Ac=.50  p_u_=0.48  p_c_=0.53 | Ac=.53  p_u_=0.11  p_c_=0.31 | Ac=.52  p_u_=0.18  p_c_=0.31 | Ac=.52  p_u_=0.18  p_c_=0.31 | Ac=.53  p_u_=0.17  p_c_=0.31 | Ac=.53  p_u_=0.10  p_c_=0.31 | Ac=.52  p_u_=0.18  p_c_=0.31 | Ac=.52  p_u_=0.15  p_c_=0.31 | Ac=.52  p_u_=0.14  p_c_=0.31 | Ac=.52  p_u_=0.19  p_c_=0.31 | Ac=.50  p_u_=0.55  p_c_=0.58 | Ac=.52  p_u_=0.16  p_c_=0.31 | Ac=.53  p_u_=0.08  p_c_=0.31 | Ac=.49  p_u_=0.58  p_c_=0.58 | Ac=.50  p_u_=0.43  p_c_=0.52 | Ac=.51  p_u_=0.27  p_c_=0.39 | Ac=.52  p_u_=0.02  p_c_=0.31 |
| ***G20-***  ***G26*** | Ac=.53  p_u_=0.30  p_c_=0.67 | Ac=.50  p_u_=0.53  p_c_=0.53 | Ac=.51  p_u_=0.40  p_c_=0.67 | Ac=.49  p_u_=0.60  p_c_=0.67 | Ac=..48  p_u_=0.66  p_c_=0.67 | Ac=.52  p_u_=0.35  p_c_=0.67 | Ac=.49  p_u_=0.55  p_c_=0.67 | Ac=.53  p_u_=0.31  p_c_=0.67 | Ac=.50  p_u_=0.47  p_c_=0.67 | Ac=.50  p_u_=0.52  p_c_=0.67 | Ac=.53  p_u_=0.27  p_c_=0.67 | Ac=.51  p_u_=0.44  p_c_=0.67 | Ac=.49  p_u_=0.56  p_c_=0.67 | Ac=.51  p_u_=0.44  p_c_=0.67 | Ac=.50  p_u_=0.50  p_c_=0.67 | Ac=.49  p_u_=0.54  p_c_=0.67 | Ac=.52  p_u_=0.36  p_c_=0.67 | Ac=.47  p_u_=0.69  p_c_=0.67 | Ac=.49  p_u_=0.57  p_c_=0.67 | Ac=.49  p_u_=0.37  p_c_=0.67 | Ac=.50  p_u_=0.51  p_c_=0.67 |
| ***G35-***  ***G37*** | Ac=.63  p_u_=0.01  p_c_=0.13 | Ac=.64  p_u_=0.01  p_c_=0.01 | Ac=.68  p_u_=4e-3  p_c_=0.09 | Ac=.56  p_u_=0.18  p_c_=0.43 | Ac=.57  p_u_=0.13  p_c_=0.43 | Ac=.55  p_u_=0.25  p_c_=0.43 | Ac=.54  p_u_=0.30  p_c_=0.43 | Ac=.55  p_u_=0.24  p_c_=0.43 | Ac=.52  p_u_=0.26  p_c_=0.45 | Ac=.55  p_u_=0.32  p_c_=0.43 | Ac=.54  p_u_=0.18  p_c_=0.43 | Ac=.52  p_u_=0.32  p_c_=0.43 | Ac=.55  p_u_=0.20  p_c_=0.43 | Ac=.57  p_u_=0.14  p_c_=0.43 | Ac=.50  p_u_=0.46  p_c_=0.43 | Ac=.58  p_u_=0.12  p_c_=0.43 | Ac=.55  p_u_=0.22  p_c_=0.43 | Ac=.54  p_u_=0.30  p_c_=0.43 | Ac=.51  p_u_=0.45  p_c_=0.46 | Ac=.51  p_u_=0.44  p_c_=0.46 | Ac=.52  p_u_=0.42  p_c_=0.46 |
| ***G40*** | Ac=.50  p_u_=0.48  p_c_=0.52 | Ac=.50  p_u_=0.49  p_c_=0.49 | Ac=.53  p_u_=0.28  p_c_=0.51 | Ac=.52  p_u_=0.33  p_c_=0.51 | Ac=.51  p_u_=0.45  p_c_=0.52 | Ac=.53  p_u_=0.22  p_c_=0.51 | Ac=.52  p_u_=0.33  p_c_=0.51 | Ac=.49  p_u_=0.59  p_c_=0.59 | Ac=.53  p_u_=0.26  p_c_=0.51 | Ac=.52  p_u_=0.31  p_c_=0.52 | Ac=.50  p_u_=0.49  p_c_=0.51 | Ac=.52  p_u_=0.33  p_c_=0.51 | Ac=.51  p_u_=0.39  p_c_=0.51 | Ac=.52  p_u_=0.33  p_c_=0.52 | Ac=.50  p_u_=0.46  p_c_=0.51 | Ac=.52  p_u_=0.37  p_c_=0.51 | Ac=.52  p_u_=0.34  p_c_=0.51 | Ac=.52  p_u_=0.31  p_c_=0.51 | Ac=.53  p_u_=0.31  p_c_=0.51 | Ac=.56  p_u_=0.13  p_c_=0.51 | Ac=.55  p_u_=0.10  p_c_=0.51 |
| ***G43*** | Ac=.51  *p_u_*=0.39  *p_c_*=0.58 | Ac=.49  p_u_=0.58  p_c_=0.58 | Ac=.51  *p_u_*=0.42  *p_c_*=0.58 | Ac=.49  *p_u_*=0.58  *p_c_*=0.58 | Ac=.51  *p_u_*=0.41  *p_c_*=0.58 | Ac=.53  *p_u_*=0.17  *p_c_*=0.58 | Ac=.50  *p_u_*=0.45  *p_c_*=0.58 | Ac=.52  *p_u_*=0.33  *p_c_*=0.58 | Ac=.50  *p_u_*=0.53  *p_c_*=0.58 | Ac=.50  *p_u_*=0.53  *p_c_*=0.58 | Ac=.51  *p_u_*=0.39  *p_c_*=0.58 | Ac=.50  *p_u_*=0.44  *p_c_*=0.58 | Ac=.50  *p_u_*=0.51  *p_c_*=0.58 | Ac=.51  *p_u_*=0.37  *p_c_*=0.58 | Ac=.49  *p_u_*=0.57  *p_c_*=0.58 | Ac=.51  *p_u_*=0.36  *p_c_*=0.58 | Ac=.50  *p_u_*=0.51  *p_c_*=0.58 | Ac=.50  *p_u_*=0.46  *p_c_*=0.58 | Ac=.52  *p_u_*=0.24  *p_c_*=0.58 | Ac=.51  *p_u_*=0.51  *p_c_*=0.58 | Ac=.55  *p_u_*=0.05  *p_c_*=0.58 |
| ***G45*** | Ac=.50  p_u_=0.52  p_c_=0.66 | Ac=.50  p_u_=0.53  p_c_=0.53 | Ac=.51  p_u_=0.40  p_c_=0.66 | Ac=.49  p_u_=0.56  p_c_=0.66 | Ac=.49  p_u_=0.63  p_c_=0.66 | Ac=.51  p_u_=0.44  p_c_=0.66 | Ac=.50  p_u_=0.54  p_c_=0.66 | Ac=.50  p_u_=0.51  p_c_=0.66 | Ac=.51  p_u_=0.41  p_c_=0.66 | Ac=.50  p_u_=0.46  p_c_=0.66 | Ac=.51  p_u_=0.42  p_c_=0.66 | Ac=.51  p_u_=0.42  p_c_=0.66 | Ac=.50  p_u_=0.49  p_c_=0.66 | Ac=.51  p_u_=0.38  p_c_=0.66 | Ac=.48  p_u_=0.64  p_c_=0.66 | Ac=.48  p_u_=0.63  p_c_=0.66 | Ac=.49  p_u_=0.55  p_c_=0.66 | Ac=.48  p_u_=0.66  p_c_=0.66 | Ac=.52  p_u_=0.31  p_c_=0.66 | Ac=.49  p_u_=0.59  p_c_=0.66 | Ac=.50  p_u_=0.5  p_c_=0.66 |
| ***G47*** | Ac=.51  p_u_=0.41  p_c_=0.48 | Ac=.50  p_u_=0.52  p_c_=0.52 | Ac=.50  p_u_=0.45  p_c_=0.48 | Ac=.53  p_u_=0.23  p_c_=0.44 | Ac=.52  p_u_=0.23  p_c_=0.44 | Ac=.53  p_u_=0.16  p_c_=0.40 | Ac=.50  p_u_=0.46  p_c_=0.48 | Ac=.53  p_u_=0.10  p_c_=0.40 | Ac=.51  p_u_=0.52  p_c_=0.48 | Ac=.51  p_u_=0.35  p_c_=0.48 | Ac=.54  p_u_=0.11  p_c_=0.40 | Ac=.50  p_u_=0.52  p_c_=0.52 | Ac=.51  p_u_=0.33  p_c_=0.48 | Ac=.53  p_u_=0.14  p_c_=0.40 | Ac=.53  p_u_=0.16  p_c_=0.40 | Ac=.54  p_u_=0.05  p_c_=0.40 | Ac=.54  p_u_=0.05  p_c_=0.40 | Ac=.51  p_u_=0.36  p_c_=0.48 | Ac=.51  p_u_=0.40  p_c_=0.48 | Ac=.52  p_u_=0.24  p_c_=0.44 | Ac=.54  p_u_=0.10  p_c_=0.40 |
| ***G55*** | Ac=.51  p_u_=0.40  p_c_=0.55 | Ac=.52  p_u_=0.36  p_c_=0.36 | Ac=.53  p_u_=0.20  p_c_=0.55 | Ac=.50  p_u_=0.36  p_c_=0.55 | Ac=.50  p_u_=0.44  p_c_=0.55 | Ac=.51  p_u_=0.38  p_c_=0.55 | Ac=.51  p_u_=0.40  p_c_=0.55 | Ac=.51  p_u_=0.42  p_c_=0.55 | Ac=.50  p_u_=0.55  p_c_=0.61 | Ac=.52  p_u_=0.31  p_c_=0.55 | Ac=.50  p_u_=0.36  p_c_=0.55 | Ac=.50  p_u_=0.45  p_c_=0.55 | Ac=.50  p_u_=0.46  p_c_=0.55 | Ac=.51  p_u_=0.37  p_c_=0.55 | Ac=.49  p_u_=0.58  p_c_=0.61 | Ac=.52  p_u_=0.32  p_c_=0.55 | Ac=.53  p_u_=0.21  p_c_=0.55 | Ac=.50  p_u_=0.47  p_c_=0.55 | Ac=.49  p_u_=0.65  p_c_=0.65 | Ac=.52  p_u_=0.30  p_c_=0.55 | Ac=.52  p_u_=0.31  p_c_=0.55 |
| ***G56*** | Ac=.51  p_u_=0.32  p_c_=0.52 | Ac=.49  p_u_=0.55  p_c_=0.55 | Ac=.51  p_u_=0.37  p_c_=0.52 | Ac=.51  p_u_=0.27  p_c_=0.52 | Ac=.52  p_u_=0.23  p_c_=0.52 | Ac=.52  p_u_=0.25  p_c_=0.52 | Ac=.50  p_u_=0.50  p_c_=0.52 | Ac=.50  p_u_=0.50  p_c_=0.52 | Ac=.50  p_u_=0.39  p_c_=0.52 | Ac=.51  p_u_=0.38  p_c_=0.52 | Ac=.50  p_u_=0.43  p_c_=0.52 | Ac=.51  p_u_=0.34  p_c_=0.52 | Ac=.50  p_u_=0.52  p_c_=0.52 | Ac=.51  p_u_=0.41  p_c_=0.52 | Ac=.50  p_u_=0.46  p_c_=0.52 | Ac=.52  p_u_=0.24  p_c_=0.52 | Ac=.51  p_u_=0.43  p_c_=0.52 | Ac=.51  p_u_=0.41  p_c_=0.52 | Ac=.51  p_u_=0.37  p_c_=0.52 | Ac=.50  p_u_=0.43  p_c_=0.52 | Ac=.54  p_u_=0.04  p_c_=0.52 |
| ***G57*** | Ac=.50  p_u_=0.49  p_c_=0.55 | Ac=.50  p_u_=0.53  p_c_=0.53 | Ac=.54  p_u_=0.24  p_c_=0.55 | Ac=.50  p_u_=0.49  p_c_=0.55 | Ac=.54  p_u_=0.24  p_c_=0.55 | Ac=.53  p_u_=0.30  p_c_=0.55 | Ac=.50  p_u_=0.47  p_c_=0.55 | Ac=.51  p_u_=0.44  p_c_=0.55 | Ac=.51  p_u_=0.44  p_c_=0.55 | Ac=.51  p_u_=0.43  p_c_=0.55 | Ac=.53  p_u_=0.26  p_c_=0.55 | Ac=.50  p_u_=0.52  p_c_=0.55 | Ac=.52  p_u_=0.37  p_c_=0.55 | Ac=.55  p_u_=0.16  p_c_=0.55 | Ac=.50  p_u_=0.52  p_c_=0.55 | Ac=.51  p_u_=0.41  p_c_=0.55 | Ac=.51  p_u_=0.39  p_c_=0.55 | Ac=.49  p_u_=0.59  p_c_=0.59 | Ac=.52  p_u_=0.34  p_c_=0.55 | Ac=.53  p_u_=0.26  p_c_=0.55 | Ac=.51  p_u_=0.45  p_c_=0.55 |
| ***G62*** | Ac=.52  p_u_=0.34  p_c_=0.67 | Ac=.50  p_u_=0.49  p_c_=0.49 | Ac=.50  p_u_=0.48  p_c_=0.67 | Ac=.52  p_u_=0.37  p_c_=0.67 | Ac=.49  p_u_=0.54  p_c_=0.67 | Ac=.51  p_u_=0.44  p_c_=0.67 | Ac=.48  p_u_=0.64  p_c_=0.67 | Ac=.48  p_u_=0.63  p_c_=0.67 | Ac=.50  p_u_=0.46  p_c_=0.67 | Ac=.47  p_u_=0.67  p_c_=0.67 | Ac=.51  p_u_=0.40  p_c_=0.67 | Ac=.51  p_u_=0.43  p_c_=0.67 | Ac=.49  p_u_=0.54  p_c_=0.67 | Ac=.49  p_u_=0.57  p_c_=0.67 | Ac=.50  p_u_=0.52  p_c_=0.67 | Ac=.51  p_u_=0.45  p_c_=0.67 | Ac=.50  p_u_=0.48  p_c_=0.67 | Ac=.48  p_u_=0.65  p_c_=0.67 | Ac=.50  p_u_=0.48  p_c_=0.67 | Ac=.50  p_u_=0.49  p_c_=0.67 | Ac=.57  p_u_=0.14  p_c_=0.67 |
| ***G80-***  ***G83*** | Ac=.49  p_u_=0.55  p_c_=0.61 | Ac=.50  p_u_=0.49  p_c_=0.49 | Ac=.54  p_u_=0.26  p_c_=0.61 | Ac=.53  p_u_=0.30  p_c_=0.61 | Ac=.51  p_u_=0.43  p_c_=0.61 | Ac=.51  p_u_=0.46  p_c_=0.61 | Ac=.55  p_u_=0.23  p_c_=0.61 | Ac=.50  p_u_=0.51  p_c_=0.61 | Ac=.50  p_u_=0.49  p_c_=0.61 | Ac=.53  p_u_=0.28  p_c_=0.61 | Ac=.51  p_u_=0.46  p_c_=0.61 | Ac=.50  p_u_=0.50  p_c_=0.61 | Ac=.55  p_u_=0.52  p_c_=0.61 | Ac=.49  p_u_=0.54  p_c_=0.61 | Ac=.50  p_u_=0.52  p_c_=0.61 | Ac=.55  p_u_=0.22  p_c_=0.61 | Ac=.52  p_u_=0.33  p_c_=0.61 | Ac=.56  p_u_=0.16  p_c_=0.61 | Ac=.48  p_u_=0.64  p_c_=0.67 | Ac=.47  p_u_=0.68  p_c_=0.68 | Ac=.52  p_u_=0.39  p_c_=0.61 |
| ***G93*** | Ac=.48  p_u_=0.61  p_c_=0.64 | Ac=.48  p_u_=0.64  p_c_=0.64 | Ac=.48  p_u_=0.61  p_c_=0.64 | Ac=.52  p_u_=0.34  p_c_=0.64 | Ac=.49  p_u_=0.57  p_c_=0.64 | Ac=.50  p_u_=0.48  p_c_=0.64 | Ac=.51  p_u_=0.46  p_c_=0.64 | Ac=.51  p_u_=0.40  p_c_=0.64 | Ac=.51  p_u_=0.43  p_c_=0.64 | Ac=.51  p_u_=0.44  p_c_=0.64 | Ac=.51  p_u_=0.46  p_c_=0.64 | Ac=.51  p_u_=0.43  p_c_=0.64 | Ac=.50  p_u_=0.47  p_c_=0.64 | Ac=.51  p_u_=0.47  p_c_=0.64 | Ac=.48  p_u_=0.62  p_c_=0.64 | Ac=.49  p_u_=0.56  p_c_=0.64 | Ac=.55  p_u_=0.37  p_c_=0.64 | Ac=.54  p_u_=0.64  p_c_=0.64 | Ac=.51  p_u_=0.53  p_c_=0.64 | Ac=.51  p_u_=0.56  p_c_=0.64 | Ac=.51  p_u_=0.39  p_c_=0.64 |
| ***Age (small)*** | Ac=.90  p_u_=7e-17  p_c_=1e-15 | | Ac=.87  p_u_=2e-12  p_c_=1e-11 | Ac=.73  p_u_=5e-5  p_c_=1e-4 | Ac=.74  p_u_=5e-5  p_c_=1e-4 | Ac=.72  p_u_=5e-4  p_c_=7e-4 | Ac=.84  p_u_=6e-11  p_c_=3e-10 | Ac=.74  p_u_=9e-4  p_c_=1e-3 | Ac=.75  p_u_=9e-5  p_c_=1e-4 | Ac=.90  p_u_=2e-15  p_c_=2e-14 | Ac=.79  p_u_=2e-3  p_c_=2e-3 | Ac=.60  p_u_=8e-2  p_c_=9e-2 | Ac=.71  p_u_=2e-4  p_c_=2e-4 | Ac=.75  p_u_=2e-5  p_c_=8e-5 | Ac=.52  p_u_=0.32  p_c_=0.34 | Ac=.74  p_u_=3e-5  p_c_=9e-5 | Ac=.73  p_u_=6e-5  p_c_=1e-4 | Ac=.52  p_u_=0.35  p_c_=0.35 | Ac=  p_u_=  p_c_= | Ac=  p_u_=  p_c_= |  |
| ***Age (large)*** | Ac=.94  p_u_=8e-68  p_c_=1e-66 | | Ac=.91  p_u_=5e-59  p_c_=4e-58 | Ac=.73  p_u_=2e-23  p_c_=3e-23 | Ac=.77  p_u_=2e-26  p_c_=3e-26 | Ac=.80  p_u_=6e-34  p_c_=1e-33 | Ac=.78  p_u_=1e-31  p_c_=2e-31 | Ac=.81  p_u_=6e-28  p_c_=9e-28 | Ac=.85  p_u_=2e-45  p_c_=7e-45 | Ac=.79  p_u_=5e-33  p_c_=1e-32 | Ac=.80  p_u_=8e-31  p_c_=1e-30 | Ac=.86  p_u_=4e-47  p_c_=2e-46 | Ac=.81  p_u_=8e-38  p_c_=3e-37 | Ac=.77  p_u_=6e-19  p_c_=7e-19 | Ac=.53  p_u_=2e-3  p_c_=2e-3 | Ac=.81  p_u_=2e-35  p_c_=6e-35 | Ac=.81  p_u_=5e-32  p_c_=9e-32 | Ac=.58  p_u_=2e-5  p_c_=2e-5 | Ac=  p_u_=  p_c_= | Ac=  p_u_=  p_c_= |  |
